# Dissecting the evolving cellular landscape of a remyelinating microenvironment

**DOI:** 10.1101/2024.12.25.630253

**Authors:** George S. Melchor, Maryna Baydyuk, Zeeba Manavi, Jingwen Hu, Jeffrey K. Huang

## Abstract

Demyelination, or the loss of myelin in the central nervous system (CNS) is a hallmark of multiple sclerosis (MS) and occurs in various forms of CNS injury and neurodegenerative diseases. The regeneration of myelin, or remyelination, occurs spontaneously following demyelination. The lysophosphatidylcholine (LPC)-induced focal demyelination model enables investigations into the mechanisms of remyelination, providing insight into the molecular basis underlying an evolving remyelinating microenvironment over a tractable time course. Here, we present a detailed analysis using high-resolution single nucleus RNA sequencing to investigate gene expression dynamics across multiple cell populations involved in the remyelination process. We examine three specific time points following focal demyelinating injury in mice, and by delineating activation states within the heterogeneous cell populations of demyelinated lesions, we highlight changes in gene expression within subclusters of each cell population from the early stages of injury response to the initiation and maintenance of remyelination. Our findings reveal how shifts in microglial, astrocytic and fibroblast activities within lesions are associated with efficient oligodendrocyte differentiation during remyelination.

## INTRODUCTION

Myelin is a lipid-rich membrane sheath generated by oligodendrocytes (OLs) in the central nervous system (CNS). OLs extend terminal processes that can engage with and enwrap axonal segments, forming myelin sheaths to facilitate rapid conduction of neuronal electrical impulses and enable metabolic coupling between OLs and neurons^1,2^. Demyelination, or the loss of myelin, is a hallmark of multiple sclerosis (MS)^3–6^ but is also prominent in other forms of CNS injury and disease including Alzheimer’s disease (AD)^7,8^, traumatic brain/spinal cord injury^9^, and stroke^10^. Demyelination has deleterious effects on neuronal health, causing impaired conduction and axonal dystrophy^3^. Further, chronically demyelinated axons, and the failure to restore myelin around them, can result in progressive neurodegeneration^11^.

In the healthy adult CNS and during the early stages of MS, myelin can naturally regenerate after demyelinating injury owing to the widespread abundance of oligodendrocyte precursor cells (OPCs) in the CNS^3^. These OPCs react to demyelinating injury by migrating to the lesion site, proliferating, and subsequently differentiating into mature OLs to generate new myelin sheaths around denuded axons^12^. The process of myelin generation through oligodendrocyte lineage cells (OLCs) after CNS injury is called remyelination. However, with chronic disease and advanced age, remyelination becomes less efficient and eventually fails, leading to progressive neurodegeneration evident in later stages of MS. Contributing factors to remyelination failure in MS include chronic inflammatory microglial activity in and around demyelinated lesions, intrinsic changes to OPCs associated with advanced age and chronic inflammation, and the persistence of cellular and membrane debris within demyelinated lesions, which hinder effective differentiation of OPCs into myelinating OLs^3,6,11^.

Remyelination, while crucially dependent on OLCs, is the result of interconnected activities involving various cell types that facilitate the process within the injury microenvironment. Several single-cell transcriptomic studies have demonstrated the heterogeneous nature of OLs, microglia, astrocytes, and vascular cells in the adult CNS. This heterogeneity in transcriptional activation states within each cell type has been suggested to play a role in either promoting demyelination or facilitating remyelination, both in mouse models of demyelination and in MS^13–17^. How the transcriptome of major cell populations within the injured microenvironment evolve over time during spontaneous remyelination remains inadequately elucidated. Therefore, it is essential to contextualize observed shifts in subpopulations or alterations in gene expression within a defined remyelination timeline to fully understand the significance of these changes in promoting remyelination. To this end, we generated a detailed transcriptomic profile of all major CNS cell types at the single nucleus level within demyelinated lesions in mice as they progress towards remyelination in the tractable lysophosphatidylcholine (LPC), or lysolecithin, model of demyelination. We observed distinct transcriptional changes and heterogeneous subsets of oligodendrocyte lineage cells (OLCs), microglia, astrocytes, fibroblasts, and endothelial cells across three post lesioned time points. Notably, the emergence of injury- and disease-associated OLC phenotypes was observed during remyelination. Microglia transitioned from early inflammatory, phagocytic states to potentially pro-regenerative profiles by later stages, while astrocytes contributed to glial scar formation and angiogenesis via extracellular matrix remodeling and displayed differential activities in early and late stage remyelination. Fibroblasts facilitated early fibrotic scar formation, and endothelial cells supported blood-brain barrier recovery. Cell populations within adjacent non-lesioned tissues exhibited transcriptional activation states not observed in naïve, unlesioned tissue, suggesting systemic responses to inflammation. These findings reveal that dynamic shifts in the activation states of microglia, astrocyte, and fibroblast subpopulations are associated with oligodendrocyte progression during remyelination, thereby highlighting the coordinated roles of CNS cells within remyelinating lesions.

## METHODS

### Mice

Mice of both sexes at 8-12 weeks old were used for each experiment. Wild-type C57BL/6J (JAX:000664) mice were obtained from Jackson Laboratories (Bar Harbor, ME), and Charles Rivers. LysMCre (JAX:004781) and B6.Cg-Gt(ROSA)26Sortm14 (JAX:007914) were also purchased from Jackson Laboratories. Allt mice were maintained on a C57BL/6 background. Mice were further maintained on a 12-hour light/dark cycle with food and water *ad libitum*. All experiments were approved by and conducted in accordance with the Institutional Animal Care and Use Committee (IACUC) at Georgetown University.

### Immunofluorescence and imaging

Mice were anaesthetized with 3% isoflurane (Pivetal) before perfusion. Mice were perfused with 4% paraformaldehyde (PFA, Sigma) in PBS. Spinal cord tissues were dissected and post-fixed in 4% PFA, followed by 30% sucrose (w/v), before freezing in Tissue-Tek optimal cutting temperature medium (Sakura) on crushed dry ice, then final storage at −80°C. Cryosectioning (Leica CM1860) was performed at 12 microns, and then sections were mounted on SuperFrost Plus slides (VWR International). Sections were dried on the slides for approximately 1 hour before storage at −80°C. For immunostaining, slides were dried at RT for 1 hour, washed, and then incubated with blocking buffer (10% donkey serum 0.25% (v/v) Triton X-100 in 1xTBS) for 1 hour at RT. Antigen retrieval pretreatment was performed by incubating slides in boiling buffer for 30 mins to optimize staining for intracellular primary antibodies. For slides utilizing mouse antibodies, an extra incubation with mouse-on-mouse blocking reagent (Vector Laboratories) was performed for 1 hour at RT. After blocking, slides were incubated with primary antibodies (**Table 1**) diluted in blocking buffer overnight at 4°C. The following day sections were washed, and then stained with secondary antibodies (Alexa Fluor 488 and 594) and Hoechst in blocking buffer for 45 minutes at RT. After final washes, slides were cover slipped with Fluoromount-G (ThermoFisher #00495802). Immunofluorescence images were collected on a Zeiss LSM 800 complete system confocal imager. Demyelinated lesion areas were located by the visualization of dense Hoechst-positive nuclei within the ventral funiculus. Quantification was conducted by 2 blinded investigators using ImageJ (manual counting).

**Table 1.**
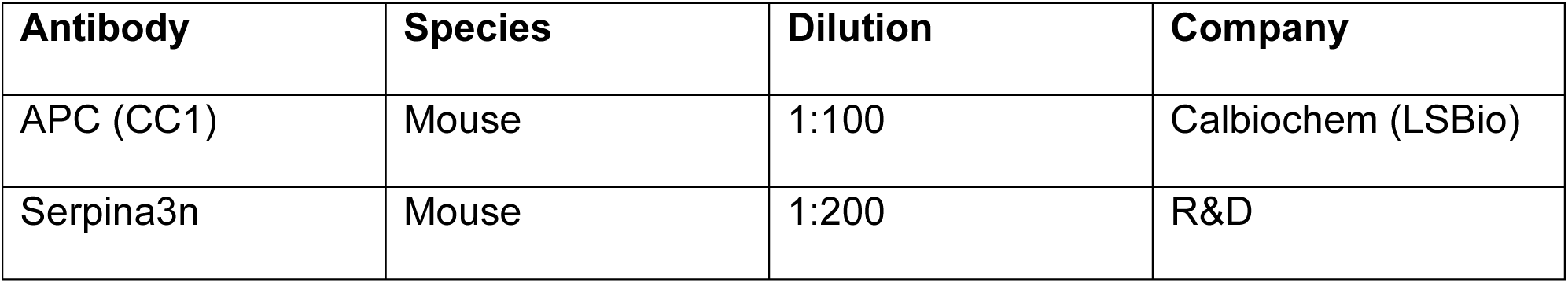
Primary antibodies used for Immunofluorescence.

### Focal spinal cord demyelination

Lysolecithin-induced focal spinal cord demyelination has been previously described^12,18^. Briefly, 1.0% lysophosphatidylcholine (lysolecithin, Sigma) is diluted in sterile PBS and then injected (1µL) into the ventral funiculus at approximately the T12 vertebra area using aseptic technique while animals are anesthetized under 3% isoflurane (Pivetal). After focal demyelination, mice were euthanized and perfused for remyelination analysis at 5 days post lesion (dpl), 10dpl, and 20dpl. Rostral, unlesioned spinal cord tissue from the same mouse was used for internal control purposes. Additionally, naïve spinal cord tissue from uninjured mice were used as external controls.

### Neutral red labeling of demyelinated lesions

To detect and microdissect lysolecithin-induced focally demyelinated lesions in spinal cords, mice were intraperitoneally injected with 0.5mL of 1% neutral red vital dye in sterile PBS two hours before PBS perfusion, as previously described^19^. Mice were anaesthetized with 3% isoflurane (Pivetal) before perfusion.

### Cell fractionation and nuclei purification

Pooled tissues were manually homogenized in Nuclei PURE lysis solution with 1M DTT (Sigma, 646563) using Dounce tissue grinders on ice. Sucrose density was prepared according to the Nuclei PURE kit and the lysates were layered over a 1.85M Sucrose Cushion in ultracentrifuge tubes (Beckman Coulter, C13926). High-speed centrifugation was carried out at 30,000 RCF (15,600 RPM) using SW28 rotor for 45 minutes at 4°C. The supernatant was discarded, and the pellet was washed twice in 1ml 2% BSA (Miltenyi Biotec) in PBS with 0.2U/µl Protector RNase inhibitor (Sigma Aldrich) followed by centrifugation at 500 RCF for 5 minutes at 4 °C. The pellet was resuspended in 100µl wash buffer and the nuclei suspension was filtered using a 40µm Flomi Cell Strainer (Sigma Aldrich). Intact nuclei morphology was verified under 40X brightfield microscope and total nuclei count was obtained using Countess II FL Automated Cell Counter.

### cDNA library preparation

The samples were processed for snRNA-seq following the protocol for *Chromium next GEM Single Cell 3’ v3.1 Dual Index* (10x Genomics). 10,000 nuclei per condition were loaded into Chromium Single Cell 3’ Chip G wells and placed in the Chromium controller for GEM generation as outlined in 10x Genomics User Guide (CG000315. Rev A) for a targeted recovery of 5,000 nuclei per condition. GEMs were incubated in the thermal cycler for cDNA barcoding, followed by GEM-RT cleanup and cDNA amplification using v3.1 Chromium GEM kit. Total cDNA concentration was measured by Qubit dsDNA HS and 25% of the total cDNA was used for 3’ gene expression library construction using Dual Index Kit TT Set A. The resulting sequencing libraries were assessed for quality using the BioAnalyzer 2100 with a High Sensitivity DNA kit (Agilent), and quantified using the KAPA Library Quantification Kit, Illumina Platforms (KAPA Biosystems) according to manufacturer’s protocol.

### Single nuclei RNA (snRNA) sequencing

Libraries were sequenced on the Illumina NovaSeq 6000 System (Illumina, USA) with a S4 FlowCell to an average depth of 400 million reads per sample at the University of Maryland (Maryland Genomics).

### Experimental groupings and statistical N for snRNAseq

Mice in the 5dpl, 10dpl, and 20dpl groups were lesioned on separate days. The surgeries were conducted by 2 of 3 experimenters for each group. Nonlesioned tissue was collected alongside lesioned tissue, providing nonlesioned tissue for each timepoint. Naïve tissue was collected with the 5dpl group. Further, 3-4 mice were pooled together for each unique sample in this study, resulting in high biological variability and a statistical N of 13. NR labeling, micro dissection of the spinal cord tissue, pooling, cell fractionation, and nuclei purification were conducted on separate days for the groups as well. cDNA library preparation was conducted separately for 5dpl and Naïve mice, while the rest of the samples were prepped on the same day. Libraries were all sequenced at the same time.

### snRNA sequencing preprocessing

The cellranger mkfastq pipeline (Cell Ranger Version 7.1.0) was used to process the sequencing run generated, described above. Following demultiplexing, the cellranger count pipeline (Cell Ranger Version 7.1.0) was used to align to the mouse transcriptome (mm10-2020-A) and quantify single nuclei gene expression. Following cellranger count, the raw matrix of each sample was run through CellBender (v0.3.0)^20^, which utilizes a deep generative learning model to reduce the amount of ambient RNA contamination that is inherently present due to cell dissociation and nucleus extraction in our sample libraries^21^. CellBender effectively constructs denoised count matrices for each sample and determines the number of nuclei present in the sample, separating out empty droplets. After CellBender implementation, a total of 85,558 nuclei across 13 samples (Naïve (2), 5dpl (3), 10dpl (2), 20dpl (3), NL5dpl (1), NL10dpl (1), NL20dpl(1)). Quality control and subsequent analyses were performed utilizing Seurat (v. 5.0.1) within R (4.4.1). Sample nuclei count matrices were filtered utilizing *ddqc*^22^, such that on a cluster-by-cluster basis nuclei with nUMI>500 and 300<nGene<3MADs were retained. Further filtration utilized universal standards of mitochondrial gene counts <1% and ribosomal gene counts <5%. Filtration resulted in a total of **71,764** nuclei across all samples and experimental conditions.

### Broad normalization and clustering

For all main figures, naïve controls and lesioned samples at all timepoints were merged and utilized in analyses. Nonlesioned controls were highlighted in Figure S6, following the same standards utilized to analyze the other experimental conditions. The Seurat (v. 5.0.1) package was used for data normalization and clustering. For clustering, data were log normalized and variance stabilized using SCTransform (v2)^23,24^. Principal component analysis was performed using the top 2,000 most variable genes and clustering analysis was performed via FindNeighbors() and FindClusters() with the top 75 PCA dimensions and a resolution of 0.4, respectively. UMAP visualizations were performed with the top 75 PCA dimensions. The count matrix (RNA assay) was also log normalized using NormalizeData() to identify cluster gene markers, expression visualizations, and for DEG calculations. The top 2,000 most variable genes were identified, and the data was scaled. Using FindAllMarkers(logfc.threshold = 1.0, min.pct =0.20) and previous knowledge of broad cell types expected in our dataset, we identified gene markers for all clusters and annotated each cluster into broad cell types: *neurons, oligodendrocyte lineage cells, astrocytes, vascular cells (inclusive of mesenchymal cells), immune cells, and Schwann cells*. The broad cell types were then subclustered and re-analyzed to investigate subpopulations across all sample conditions. Most visualizations utilize a combination of scCustomize (v2.1.2, DOI: 10.5281/zenodo.5706430), Seurat (v5.0.1), and ggplot2 (v3.5.1).

### Subclustering

Major glial cell populations including oligodendrocyte lineage cells (OLCs), immune cells, astrocytes, and vascular/mesenchymal cells were subclustered and analyzed. For each subclustered broad cell type, after the initial subset was conducted, nuclei were cleaned up manually post-clustering based on the expression of multiple population marker genes and low quality indicating they were likely doublets. This manual cleanup resulted in a smaller number of nuclei being analyzed for every subset than was originally annotated (**Table 2**). After this clean up, data were normalized and variance stabilized with SCTransform (v2) PCA was performed, and clustering analysis was conducted following the same standard pipeline (using the top 65 dimensions) as in the broad normalization and clustering. The count matrix (RNA assay) also followed the same standard pipeline to enable cluster marker gene identification, for plotting of expression values, and for DEG calculations.

**Table 2.** SubCluster nuclei manual cleanup.

| Cell type | Starting Nuclei | Doublets and LowQ | % Doublets and LowQ | Nuclei Analyzed |
| --- | --- | --- | --- | --- |
| OLC | 20,719 | 293 | 1.41% | 20,426 |
| Immune | 3,926 | 351 | 8.94% | 3,575 |
| Astrocyte | 8,017 | 800 | 9.98% | 7,217 |
| Vascular | 6,294 | 807 | 12.82% | 5,487 |

### Differential gene expression (DEG) testing and identification

Differential expression of genes in pairwise comparisons between experimental conditions and annotated subclusters was conducted using FindMarkers() from Seurat, with the MAST test^25^ using parameters: logfc.threshold() = 1.0 and min.pct = 0.2. The adjusted p-value was calculated by the function using Bonferroni correction. DEGs were filtered such that only those with an adjusted p-value < 0.05 were considered significant.

### Gene set enrichment analysis

DEG lists were compiled from pairwise comparisons in this study and used as input to Metascape (v3.5.20240901, https://metascape.org)^26^, along with background gene lists calculated for each subclustered broad cell type with a threshold for mean expression set at 0.1. Express analysis with default settings was used for determining gene set enrichment.

### Volcano plots, Venn diagrams, and correlation heatmaps

Volcano plots were created using EnhancedVolcano (v1.22.0), Venn diagrams were created using ggVennDiagram (v1.5.2), and Pearson correlation heatmaps were generated with SCpubr (v2.0.2)^27^.

### Scaled heatmaps

Scaled heatmaps were generated with SCpubr (v2.0.2). Z score transformed scaled data calculated from normalization and scaling methods (*see Broad normalization and clustering)* was used as input for heatmaps along with gene sets of interest. All individual samples pertaining to the comparisons made in heatmaps are included. Further, compared groups in heatmaps were randomly downsampled to avoid bias in the visual representation of the data.

### Gene set score calculations

Gene set scores were calculated by the aggregate normalized sum expression of genes present in each gene set per nuclei barcode. The white matter associated microglia (WAM) score was based off of 100 WAM genes identified in aged and AD mice^17,28^. The necroptosis gene set score was based off of 16 genes correlated to necroptosis following LPC-mediated demyelination in the CNS^29^. The SenMayo score was determined by using 113 mouse-specific senescence-associated genes compiled from extensive review and comparative analysis of multiple age-related datasets^30^. The disease associated microglia (DAM) score was determined by using the top 71 DAM genes identified in the 5xFAD mouse model^17,31^. Complete gene set lists can be found in Table S3. Gene set scores were visualized using the FeaturePlot_scCustom() function.

### Frequency bar plots, stacked DEG bar plots, trendlines, and gene set enrichment bar plots

Cluster relative frequency bar plots were generated with SCpubr (v2.0.2). Trendlines, stacked DEG bar plots, and gene set enrichment pathway charts were created using customized ggplot2 code. For normalized gene expression trendlines, mean expression was calculated for selected genes and then normalized by subtracting mean expression of each respective gene from the naïve condition. The theme aesthetics for trendlines and gene set enrichment bar plots utilized a custom plot_theme() function borrowed from the broader R community. Confidence intervals (95%) were calculated and were plotted using the geom_ribbon() function.

### Quantification and statistical analysis

All statistical tests used for comparisons are included in figure legends. All statistical analyses for validation IF were performed using Prism 10.0 software with students T-test, p value < 0.05, *. P values in snRNA-seq analysis for cluster markers and DEGs were adjusted for multiple comparisons based on Bonferroni correction.

### Illustrations

All illustrations that support the presented data were generated in BioRender.

### Data and code availability

Single-nucleus RNA sequencing transcriptomic data is deposited in the Gene Expression Omnibus (GEO) database at accession number GSE278643. and will be publicly available at the date of publication. Original code developed in the processing of the data in this study will be deposited in Github under a publicly available link at the time of publication. Any additional information required to reanalyze the data is available from Jeffrey K. Huang, the contact author, upon request.

### Supplemental Information

Table S1. Excel file containing cluster markers for annotated populations.

Table S2. Excel file containing annotated cluster stats.

Table S3. Excel file containing external gene lists utilized for gene set score comparisons.

## RESULTS

### Changes in the proportion of cellular compositions and gene expression within lesions during remyelination

LPC-induced demyelination creates a contained, focal lesion which follows a reproducible and tractable timeline of remyelination that involves oligodendrocyte lineage cell (OLC) progression and immune and glial cell activation^12,18^. To define the transcriptome of the lesion microenvironment during remyelination, we induced focal demyelination by injecting 0.1% LPC into the ventral spinal cord white matter of 8–12-week-old female and male mice. Focal lesions were then identified using neutral red labeling^19^. Next, we performed single nucleus RNA sequencing (snRNAseq) on dissected spinal cord white matter tissues from (1) naïve, unlesioned control, (2) lesions at 5 days post lesion (dpl), corresponding to early injury responses and OPC proliferation, (3) lesions at 10dpl, corresponding to OPC differentiation, (4) lesions at 20dpl, corresponding to late stages of remyelination, and (5) adjacent, nonlesioned spinal cord tissue from lesioned animals at all 3 timepoints (**Figure 1A**).

**Figure 1.**
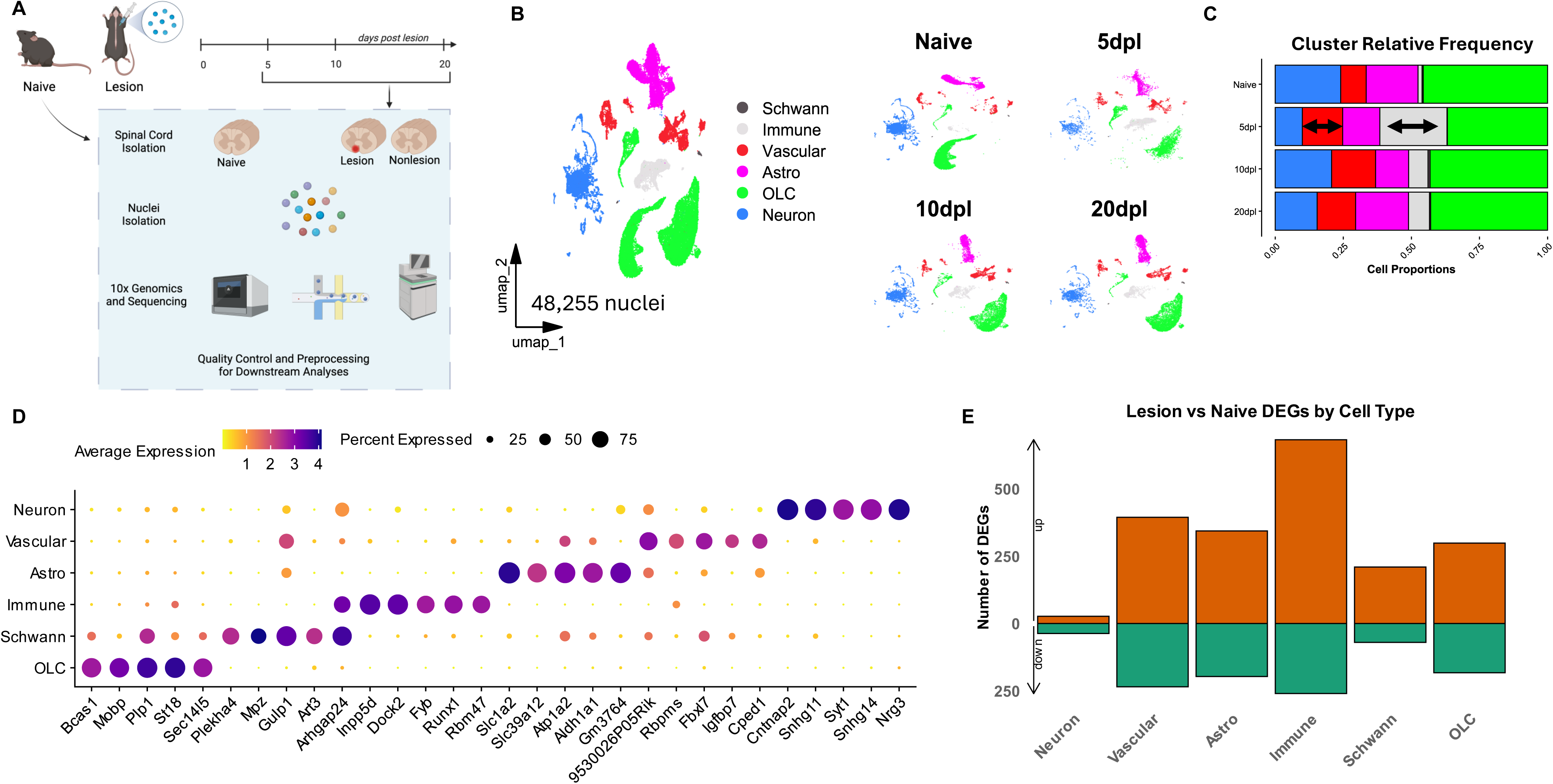
Multiple glial cell populations are profoundly impacted by LPC-mediated focal demyelination and subsequent remyelination. **A)** Schematic diagram of the experimental design and overall strategy. Created using BioRender.com. **B)** UMAP plot of 48,255 broadly annotated lesion and naïve nuclei clusters, including neurons (Neuron), oligodendrocyte lineage cells (OLCs), astrocytes (Astro), vascular and mesenchymal cells (Vascular), immune cells (Immune), and Schwann cells (Schwann, *left*), split by timepoint (*right*).**C)** Bar plot showing the relative frequencies of broadly annotated cell populations split by timepoint. Black double-sided arrows highlight vascular and immune cell increases at 5dpl. See Table S2 for the number of nuclei and relative frequencies of all annotated clusters. **D)** Dot plot showcasing top marker genes used to annotate broad cell populations. See Table S1 for a complete list of cluster markers. **E)** Stacked bar plot indicating the number of upregulated (*top, orange*) and downregulated (*bottom, green*) genes calculated in pairwise comparisons between Naïve and total Lesion samples for each broadly annotated cell population. Log2FC >1.0, min. pct = 0.2, adjusted p-value <0.05, MAST, Bonferroni correction.

Following quality control (*see Methods),* we analyzed a total of **48,255** nuclei from lesioned samples and naïve controls (**Figures 1B, S1A-S1C**). The merged nuclei were clustered and broadly annotated into 6 major CNS cell types based on known and calculated enriched transcriptional signatures (**Figures 1B**, **1C and 1D; Table S1**), which included *Bcas1*, *Mobp*, and *Plp1* for OLCs, *Mpz* and *Plekha4* for Schwann cells, *Inpp5d*, *Dock2*, and *Runx1* for immune cells, *Slc1a2*, *Slc39a12*, and *Aldh1a1* for astrocytes, *Cped1*, *Rbpms*, and *Igfbp7* for vascular and mesenchymal cells, and *Cntnap2*, *Snhg11*, and *Syt1* for neurons. We found that the frequency of most major cell populations examined were affected over the course of remyelination (**Figures 1B and 1C; Table S2**). Shortly following demyelination, at 5dpl, we observed a reduction in neurons, astrocytes, and OLCs, all of which increased towards control levels at later timepoints 10 and 20dpl. A substantial increase in immune and vascular nuclei was also observed at 5dpl. Notably, these cell populations decreased by 20dpl, yet remained elevated compared to naïve controls (**Figures 1B and 1C; Table S2**). Along with the alterations in cell type frequency, OLCs, astrocytes, vascular, and immune cells all showed significantly increased differentially expressed genes (DEGs) in the lesion (**Figure 1E**). Only a small number of DEGs were identified in neurons, and although Schwann cells showed a large number of DEGs, they represented only a minor proportion at any time point (**Figures 1B and 1C; Table S2**). Together, these results suggest LPC-mediated focal remyelination primarily affects glial and vascular populations in the mouse spinal cord.

### OPC differentiation initiates early after demyelination, leading to heterogeneous subsets of mature oligodendrocytes during remyelination

To assess the effects of the remyelination process following LPC-mediated demyelination on OLC transcriptomes, we subclustered OLCs, the largest relative cell proportion of all of our samples (**Figure 1C; Table S2**).We observed fewer nuclei in lesion timepoints compared to naive (**Figure S2A**), and major alterations in transcriptional programs (**Figure S2B**). We further annotated OLC subclusters based on previously defined^17,32–35^ and calculated gene markers (**Figures 2A and S2C; Table S1**). In naïve samples, most nuclei were mature OL (MOL) subpopulations: MOL2/3 or MOL5/6, distinguished by the expression of *Cntn3*, *Tspan9*, *Enpp6* and *Ptgds*, *Il33*, and *Opalin*, respectively, and MOL1-6 exhibiting a mixture of both markers (**Figures 2B and S2C, top; Table S2**). Oligodendrocyte precursor cells (OPCs), demarcated by *Pdgfra*, *Cspg4*, and *Cspg5* and myelin forming oligodendrocytes (MFOLs) demarcated by *Synpr*, *Kit*, and *Man1a*, constituted the majority of the remaining naïve nuclei (**Figures 2B and S2C, bottom; Table S2**). At 5dpl, MOL populations decrease dramatically (**Figures 2A and 2B; Table S2**), consistent with previous LPC-mediated demyelination studies^19,36,37^. By 10 and 20dpl, MOLs steadily increase to near homeostatic frequency levels (**Figures 2A and 2B; Table S2**), suggesting successful OLC lineage progression and remyelination.

**Figure 2.**
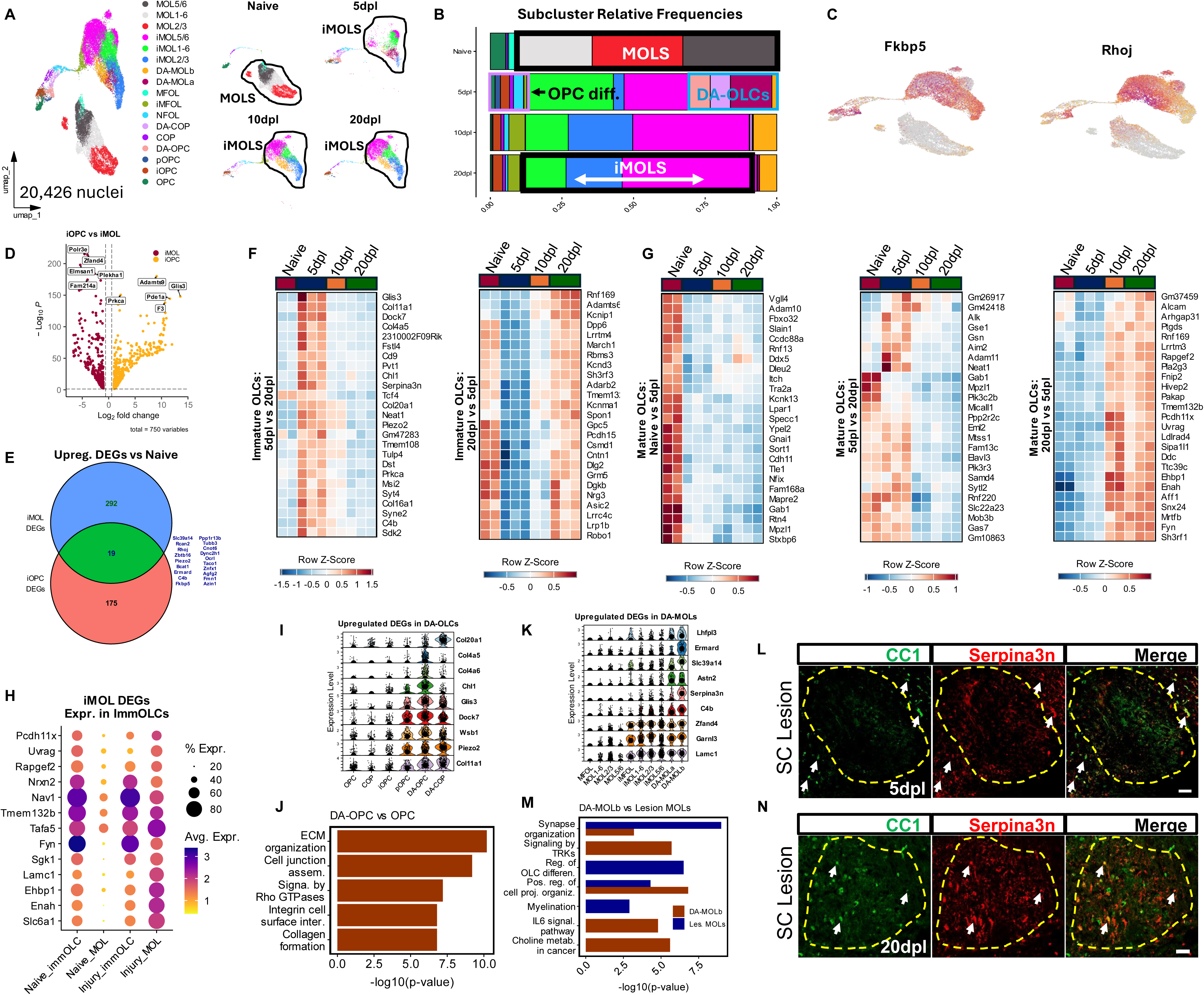
Multiple injury-associated, activated phenotypes define OLC subpopulations throughout remyelination. **A)** UMAP plot of 20,426 annotated oligodendrocyte lineage cell nuclei (*left*), split by timepoint (*right*). **B)** Bar plot showing the relative frequencies of annotated OLC subpopulations in each experimental timepoint. See Table S2 for the number of nuclei and relative frequencies of all annotated clusters. **C)** Feature plot showcasing the upregulated expression of injury-associated genes *Fkbp5* and *Rhoj* in lesion OLCs. **D)** Volcano plots showcasing DEGs calculated from a pairwise comparison of injury associated immature OLCs (iOPC, pOPC, DA-OPC, DA-COP) and mature OLCs (iMFOL, iMOL1-6, iMOL2/3, iMOL5/6, DA-MOLa, DA-MOLb). iOPC in gold, iMOL in dark red. **E)** Venn diagram showcasing the overlap (19 genes) in upregulated DEGs in injury-associated subclusters calculated from pairwise comparisons of injury associated immature OLCs vs OLCs (175 unique) and iMOLs vs MOLs (292 unique). The overlapping genes are shown. **F)** Scaled heatmaps showing differential gene expression patterns in immature OLCs (OPCs, iOPCs, pOPCs, DA-OPCs, COPs, NFOLs) across all samples. Selected genes were calculated from pairwise comparisons between immature OLCs at 5dpl vs 20dpl, showing 5dpl upregulation (*left*), and 20dpl upregulation (*right*). Genes were further selected by top ranking based on adjusted p value. **G)** Scaled heatmaps showing differential gene expression patterns in mature OLCs (MFOL, iMFOL, MOL1-6, MOL2/3, MOL5/6, iMOL1-6, iMOL2/3, iMOL5/6, DA-MOLa, DA-MOLb) across all samples. Selected upregulated genes were calculated from pairwise comparisons between MOLs at Naïve vs 5dpl (*left*), 5dpl vs 20dpl (*center*), and 20dpl vs 5dpl (*right*). Genes were further selected by top ranking based on adjusted p value. **H)** Dot plot of injury associated and naïve associated immature and mature OLC clusters showcasing several activation genes upregulated in iMOLs that are prominently expressed in immature clusters, calculated form pairwise comparisons between iMOLs and MOLs. **I)** Stacked violin plot showcasing several activation genes calculated from pairwise comparisons between DA-OPCs and OPCs, upregulated in DA immature OLC subclusters, including proliferating OPCs (pOPCs). Central dots represent the median gene expression. **J)** Bar plots showcasing gene set enrichment terms associated with calculated upregulated DEGs in DA-OPCs vs OPCs, organized by −log10(p-value). Gene set enrichment was conducted using Metascape. **K)** Stacked violin plot showcasing several activation genes calculated from pairwise comparisons between DA-MOLs and MOLs, upregulated in many injury associated MOL subclusters, including iMFOLs. Central dots represent the median gene expression. **L)** Representative immunostaining showing significant overlap between Serpina3n and CC1+ MOLs outside of demyelinated lesions at 5dpl. CC1 in green (*left*), Serpina3n in red (*center*), merge (*right*). Dashed outline denotes lesion area, scale bar = 50µm, arrows denote colocalization. **M)** Bar plots showcasing gene set enrichment terms associated with calculated upregulated DEGs in DA-MOLs vs lesion iMOLs, organized by −log10(p-value). Bars represent the −log10(p-value) for upregulated DEGs in DA-MOLs (*brown*) and upregulated DEGs in iMOLs (*blue*). Gene set enrichment was conducted using Metascape. **N)** Representative immunostaining showing significant overlap between Serpina3n and CC1+ MOLs inside LPC-mediated remyelinating lesions at 20dpl. CC1 in green (*left*), Serpina3n in red (*center*), merge (*right*). Dashed outline denotes lesion area, scale bar = 50µm, arrows denote colocalization.

Following LPC-mediated demyelination OPCs migrate to the lesion site and generate new MOLs to facilitate remyelination^12^. In our dataset, we see evidence for OPC differentiation as early as 5dpl (**Figures 2A and 2B; Table S2**). Committed to differentiation OPCs (COPs) and newly formed oligodendrocytes (NFOLs), identified by *Bmp4*, *Bcas1os2*, *Chd3* and *Rasgef1b*, *Tcf7l1*, and *Rras2*, respectively, are subsets of OLCs that represent the earliest stages of OPC differentiation^32^. COPs and NFOLs emerge and peak at 5dpl (**Figures 2A and 2B; Table S2**), suggesting most OPCs that will differentiate, initiate this process early and perhaps within a defined timeframe. At 10 and 20dpl, we observed a significant increase in MFOLs (**Figures 2A and 2B; Table S2**), which are late stage differentiating OLCs^32^. MFOLs bridge the transcriptional gap between OPCs and MOLs by expressing both immature genes (*Kirrel3*, *Pcdh7*) and myelin-related genes (*Mobp*, *Mog*) while lacking expression of mature markers, such as *Spock3*, *Anln*, and *Hapln2* (**Figure S2D).** Moreover, MFOLs express markers associated with MOL5/6, including, *Clmn*, *Ptgds*, and *Opalin* (**Figure S2D**), suggesting MFOLs may differentiate into MOL5/6 (**Figure S2E**). LPC-mediated focal demyelination kills OLCs in a well-defined lesion site, while predominantly sparing surrounding OLCs^38–40^. At 5dpl, both MOL2/3 and MOL5/6 populations are severely diminished (**Figures 2A and 2B; Table S2**), suggesting they were depleted within the lesion. By 10dpl, MOL2/3 and MOL5/6 subpopulations increase indicating their repopulation within the lesion (**Figures 2A and 2B; Table S2**), with MOL5/6 reaching levels higher than those seen in naïve samples (**Figure 2B; Table S2**). This observation further suggests MFOL may preferentially transition into MOL5/6 during remyelination, a phenomenon that was previously observed in spinal cord injury^41^.

### Remyelinating lesions consist of injury-associated and disease-associated OLC subpopulations

In addition to and concurrent with canonical lineage progression, we observed the emergence of an injury-associated OLC (iOLC) phenotype at 5dpl that persisted through 20dpl (**Figures 2A and 2B; Table S2).** iOLCs exhibited an upregulation of stress-related *Fkbp5* and actin molecule *Rhoj* (**Figure 2C**), suggesting a generalized response to injury across OLCs. Moreover, we found that immature (all OPC, COP, and NFOL subclusters) and mature OLCs (all MOL and MFOL subclusters) exhibited vastly different injury-associated profiles (**Figures 2D and 2E**). In immature OLCs, the injury associated phenotype predominated at 5dpl (**Figure 2F, left**). By 20dpl, immature OLCs upregulate expression of several genes found in homeostatic, naïve samples (**Figure 2F, right**), suggesting that by 20dopl immature OLCs may return to homeostasis. Conversely, the injury associated phenotype in MOLs (including MFOL and iMFOLs) not only spanned across all lesion and nonlesion timepoints (**Figures 2A**, **2B and S6I**), but actually was most prominent at later timepoints 10 and 20dpl (**Figure 2G, right**). In fact, injury-associated MOLs (iMOLs) were so similarly and strongly activated that they clustered separately from the homeostatic, naïve samples (**Figures. 2A**, **2B and S6I; Table S2**). The observed clustering occurred despite iMOLs exhibiting no difference in traditional myelin-related genes like *Mbp*, *Qk*, and *Mobp* (**Figure S2F**), instead highlighting differences in other myelin-related genes such as *Mpzl1*, *Rtn4*, *Gab1*, and *Mapre2* (**Figures 2G, left and S2F**). Moreover, we observed that several upregulated DEGs in iMOLs were highly expressed in immature OLC populations (**Figure 2H**). Together these data suggest that injury-associated, remyelinating MOLs at 10 and 20dpl may be partially transcriptionally immature or have yet to return to a homeostatic state.

A subset of iOLCs further exhibited a “disease-associated” state with high expression of the proteolytic inhibitor *Serpina3n* and the classical complement factor *C4b* (**Figures S2B, S2C, and S2G**), seen in OLCs across multiple animal models and in human neurodegenerative disease, including MS^17,34,35,42,43^. Disease-associated (DA) immature OLCs, including DA-OPCs and DA-COPs, emerged by 5dpl and appear to be predominantly transient, as they were not observable by 10dpl (**Figures 2A and 2B; Table S2**). DA-OPCs and DA-COPs upregulated similar activation genes including transcription factor and thyroid hormone activating *Glis3*, mechanosensitive ion *Piezo2*, thyroid activating and senescence factor *Wsb1*, and several collagen genes (**Figure 2I**). DA-OPCs were more activated than DA-COPs, exhibiting higher differential gene expression, and almost all upregulated DEGs calculated in DA-COPs were shared with DA-OPCs (**Figure S2H**). Moreover, DA-OPCs exhibited an enrichment of DEGs involved in collagen formation and extracellular matrix organization (**Figure 2J**), suggesting a contribution to glial scar formation following demyelinating injury. Additionally, despite no injury-associated cluster for NFOLs, NFOLs from lesioned samples exhibited upregulation of DEGs found in immature DA populations (**Figure S2I**), suggesting that the injury-associated and DA phenotype continually define differentiating OLC populations in lesions throughout remyelination. In contrast to immature DA-OLCs, DA-MOLs appeared by 5dpl and remained at elevated levels at 20dpl, our latest remyelination timepoint analyzed (**Figures 2A and 2B, Table S2**). DA-MOLs were observed in two major clusters, DA-MOLa associated with 5dpl and DA-MOLb associated with 10 and 20dpl (**Figures 2A and 2B; Table S2**). We found DA-MOLs to be the most activated MOLs within lesions, as they upregulated several activation genes shared with other iMOLs including *Zfand4* and *Rhoj*, as well as more DA-MOL specific genes including *Serpina3n*, *Astn2*, and *Slc39a14* (**Figure 2K**).

DA-MOLs are suggested to emerge as a response to injury by preexisting MOLs and/or through the differentiation of DA-OPCs^43^. Since mature OLs are sparsely observed in lesions at 5dpl^19,36,37^, DA-MOLa likely represents MOLs near or just outside of the lesion site (**Figure 2L**), that were activated towards a disease-associated state in the vicinity of demyelination. Notably, all DA-MOLa express transcripts found in MOL2/3 (**Figures 2A and S2I**), which has been shown to be the most abundant MOL subgroup in the adult homeostatic mouse spinal cord white matter^33^. Moreover, DA-MOLb, present at 10 and 20dpl, consist of transcripts found in both MOL2/3 and MOL5/6 (**Figures 2A and S2J**) and may represent newly generated mature OLs inside the lesion itself. Interestingly, considering the pool of MOLs within lesions, DA-MOLb exhibit upregulation of activation pathways unrelated to OLC differentiation or myelination (**Figure 2M**), suggesting that DA-MOLb may not be directly involved in the remyelination of axons or may be responding to the inflammatory environment. Further, immunostaining analysis for Serpina3n+/CC1+ expression revealed a large proportion of CC1+ MOLs inside the lesion are Serpina3n+ (**Figure 2N**), indicating that many DA-MOLs occupy the lesion at 20dpl. Top DEGs between DA-MOLb and DA-MOLa (**Figure S2K**) reveal distinctions in gene expression when compared amongst activated subclusters (**Figure S2L**) and experimental timepoints (**Figure S2M**), where we found that DA-MOLa also exhibit an enrichment of naïve MOL genes *Mob3b*, *Mpzl1*, and *Mapre2*. While higher expression of naïve-related MOL genes at 5dpl requires further investigation, these data suggest that DA-MOLb and by extension 10 and 20dpl iMOLs are transcriptionally distinct from naïve MOLs.

By 20dpl, despite the increase in MOLs inside the lesion (**Figures 2A**, **2B, and 2N**) which is an indicator of spontaneous remyelination^18^, we observed high levels of lingering stress and injury related genes such as *Fkbp5*, *C4b*, and *Serpina3n*, immature-related genes such as *Fyn*, *Sgk1*, and *Enah*, and the reduction in naïve-MOL genes such as *Mpzl1*, *Rtn4*, and *Mapre2* (**Figures 2G**, **2N, and S2F**). Taken together, these data suggest that lesion-related remyelinating MOLs are highly active through 20dpl, which may interfere with their ability to fully mature. The sustained transcriptional alterations observed could explain why regenerated myelin is often thinner than homeostatic myelin and could serve as targets for therapeutic research to enhance remyelination.

### Remyelination is associated with a shift in microglial activity towards homeostasis

Immune cells, including microglia and macrophages, respond to demyelination by expanding (**Figures 1B**, **1C**, **3A and S3A**) and initializing activation profiles that can both exacerbate neuroinflammatory damage caused by demyelination and support the remyelination process^13,29,44^. As remyelination progresses after LPC-mediated demyelination, we observed significant transcriptional changes in immune cells over time. At 5dpl, we observed the largest number of DEGs when compared to naïve controls (**Figure S3B**). Classic microglial inflammation-related genes such as *Iqgap1*, *Arhgap24*, and *Cd300lf* were upregulated at this stage (**Figure S3C, right**), while homeostatic microglial genes such as *Atp8a2*, *Siglech*, and *Selplg* were all significantly downregulated (**Figure S3C, left**). Notably, a significant number of DEGs remain detectable at 20dpl compared to naïve controls (**Figure S3B**). Further analysis of DEGs revealed two coexisting processes at this late stage in remyelination: an upregulation of a distinct set of activation markers including *Apoe*, *Fkbp5*, and *Slc1a3*, and an increase in homeostatic gene expression (**Figures S3C, left, and S3D**). These data suggest that immune cells exhibit differential activation states throughout remyelination, with a return towards homeostatic gene expression characterizing later stages of remyelination.

**Figure 3.**
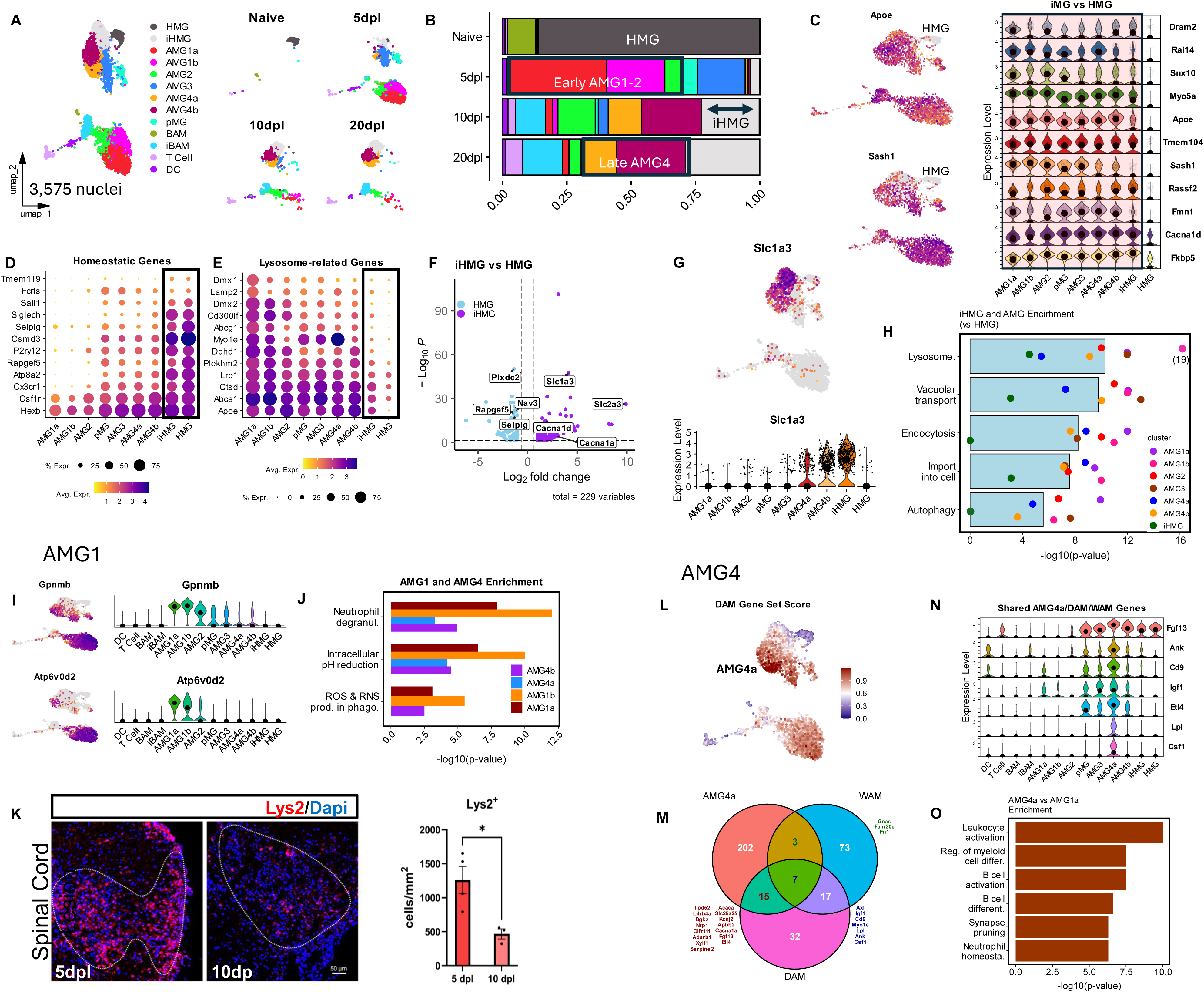
Microglial cell homeostasis and activation shifts are indicative of late stage remyelination. **A)** UMAP plot of 3,575 annotated immune nuclei (*left*), split by timepoint (*right*). **B)** Bar plot showing the relative frequencies of annotated immune subpopulations in each experimental timepoint. **C)** Feature plot (*left*) and stacked violin plot (*right*) showcasing the expression of several pan injury-associated upregulated activation genes, including *Apoe* and *Sash1*, across microglia subclusters. Central dot in stacked violin plot represents the median expression. **D)** Dot plot showing the average expression of several homeostatic genes amongst microglia subclusters. Black rectangular selection highlights high homeostatic gene expression in iHMG and HMG. **E)** Dot plot showing the average expression of several activated lysosomal genes amongst microglia subclusters. Black rectangular selection highlights low lysosomal gene expression in iHMG and HMG. **F)** Volcano plot showcasing DEGs calculated from a pairwise comparison of iHMG (*purple*) vs HMG (*blue*). Log2FC >1.0, min. pct = 0.2, adjusted p-value <0.05, MAST, Bonferroni correction. **G)** Feature plot (*top*) and violin plot (*bottom*) highlighting the expression of *Slc1a3* amongst annotated subclusters. Central dot in violin plot represents median expression. **H)** Bar plot showcasing gene set enrichment terms associated with calculated upregulated DEGs between lesion iHMG and AMGs vs naïve HMG controls, organized by −log10(p-value). Bars represent the mean of all 7 activated subclusters. Individual points represent the −log10(p-value) for AMG1a (*purple*), AMG1b (*pink*), AMG2 (*red*), AMG3 (*brown*), AMG4a (*blue*), AMG4b (*gold*), and iHMG (*green*) in each enrichment term. Gene set enrichment was conducted using Metascape. **I)** Feature plot (*left*) and violin plot (*right*) showcasing the expression of activated *Gpnmb* and *Atp6v0d2* amongst annotated subclusters. Central dot in violin plot represents median expression. **J)** Bar plot showcasing gene set enrichment terms associated with calculated upregulated DEGs shared between AMG1 and AMG4 vs naïve HMG controls, organized by −log10(p-value). Bars represent the −log10(p-value) for upregulated DEGs in AMG1a (*darkred*), AMG1b (*orange*), AMG4a(*blue*), and AMG4b(*purple*). Gene set enrichment was conducted using Metascape. **K)** Representative immunostaining showing the amount of Lys2+ cells in lesions, Lys2 tdtomato reporter in red, dapi in blue, dashed outline denotes lesion area, scale bar = 50µm, p-value <0.05, *. (*left*), and quantification of Lys2+ reporter cells indicating a significant reduction by 10dpl, n=3-4 mice (*right*). **L)** Feature plot highlighting calculated DAM gene set scores on the projected immune UMAP. AMG4a exhibited the highest average levels of DAM gene set expression. See Table S3 for the complete gene set lists used. **M)** Venn diagram showing the overlap of AMG4a cluster markers with WAM and DAM gene sets. The genes shared between AMG4a and WAM (*green*) AMG4a and DAM (*red*), and shared between all groups (*blue*) are shown. FindAllMarkers() was used to determine cluster markers. See Table S3 for the complete gene set lists used. **N)** Violin plot highlighting key late-stage DAM gene expression in immune annotated clusters. AMG4a exhibited the highest expression of several DAM genes. Central dot represents median expression. **O)** Bar plot showing selected gene enrichment terms associated with upregulated DEGs in AMG4a when compared to AMG1a, highlighting activation differences. Gene set enrichment was conducted using Metascape.

To further characterize these transcriptional alterations, we applied known^45,46^ and calculated marker genes to subcluster and annotate immune nuclei (**Figures 3A**, **3B and S3E; Table S1**). In the naïve condition, nearly all nuclei corresponded to homeostatic microglia (HMG) and border associated macrophages (BAMs) (**Figures 3A and 3B; Table S2**). After demyelination (at 5, 10, and 20dpl), we observed an increase in T cells and dendritic cells (DCs), and the emergence of proliferating microglia (pMG), injury-associated homeostatic microglia (iHMG), and several activated microglia (AMG) subpopulations, which we defined as AMG1a/b, AMG2, AMG3, and AMG4a/b. (**Figures 3A**, **3B, and S3E; Table S2**). Both iHMGs and AMG subclusters exhibited high expression of genes such as cholesterol transporter *Apoe*, TLR4-related *Sash1*, and stress-related *Fkbp5* (**Figure 3C**). Unlike AMGs, iHMGs expressed high levels of homeostatic genes (**Figures 3D and S3E; Table S1**) and had much lower expression of lysosome-related genes (**Figure 3E**), indicative of reduced phagocytic functionality. Moreover, iHMGs (and AMG4a/b) expressed high levels of *Slc1a3* (**Figures 3F**, **3G, and S3E**), a glutamate transporter previously reported in microglia following injury^47,48^, linked to extracellular glutamate clearance^49^. iHMG and AMG4a/b also significantly increase as remyelination progresses to 20dpl (**Figures 3A and 3B; Table S2**). Taken together, these data suggest iHMG are both increasingly homeostatic yet also retain levels of an injury-associated phenotype that could be beneficial to the progression of remyelination, perhaps through glutamate clearance.

Among the AMG subclusters, AMG3 and AMG4a/b exhibited elevated expression of several genes in common with HMGs such as *Cx3cr1* and *Plxdc2*, whereas AMG1a/b and AMG2 exhibited macrophage-like features, as denoted by the expression of *F13a1*^13^ and *Adgre5*^50^. (**Figures S3F, S3G, and S3H**). Notably, while all AMGs exhibit enrichment of differentially expressed genes associated with lysosome-related pathways, AMG1a/b, consistently showed the highest enrichment with AMG4a/b showing lower enrichment (**Figure 3H**). Since AMG1a/b subclusters are almost exclusively found in lesions at 5dpl (**Figures 3A**, **3B, and S6M; Table S2**), these microglial subsets may play a crucial role in the clearance of myelin debris following demyelination^51,52^. AMG1a/b subclusters also displayed high expression of *Gpnmb* (**Figures 3I and S3E**), a glycoprotein associated with Alzheimer’s (AD) pathology and myelin clearance in MS lesions^31,53,54^, and *Atp6v0d2* (**Figures 3I and S3E**), a myeloid-specific subunit of the lysosomal V-ATPase complex associated with autophagy and phagocytosis in injury and AD^31,55–57^.

AMG1a/b subclusters were further strongly associated with several classical activation pathways such as ROS & RNS production, neutrophil degranulation, and intracellular pH reduction (**Figure 3J**), indicating high levels of inflammatory activity. By 10dpl, the number of observed AMG1a/b declined significantly (**Figures 3A and 3B; Table S2**), possibly due to necroptosis or senescent cell clearance (**Figure S3I**)^29,58^. The reduction of AMG1a/b was confirmed via expression of *Lyz2* (**Figure S3J**) by immunofluorescence analysis of lesions in Lys2cre:Tdtomato mice, which labels activated myeloid cells^37^. Analysis revealed a significant reduction in TdTomato+ cells from 5 to 10dpl (**Figure 3K**), indicating a reduction in *Lyz2*+ AMG1 as remyelination progresses (**Figure S3K**). Further, AMG1a showed elevated expression of several activated myeloid markers (**Figures 3C and S3E; Table S1**), including *Lgals3*, *Spp1*, and *Lipa* (**Figures S3L, S3M, and S3N**), resembling white matter associated microglia (WAM; **Table S3**) implicated in myelin debris clearance in neurodegenerative disease and aging^28^. Altogether our observations indicate AMG1a/b subclusters are highly inflammatory and likely play a critical role in early myelin debris clearance.

In contrast to AMG1a/b, AMG4a/b subclusters were prominent at later remyelination stages (10 and 20dpl) (**Figures 3A and 3B; Table S2**). Additionally, unlike macrophage-like microglia (AMG1a/b and AMG2), AMG4a/b displayed high expression of homeostatic genes (**Figure 3D**) suggesting these subclusters are less activated. We further compared AMG4a, AMG4b, and HMGs (as positive controls) against macrophage-like AMG1a. We found that AMG4a/b DEGs were enriched in pathways related to HMG homeostatic functions such as efferocytosis and hemostasis (**Figure S3O**). While all AMG subclusters exhibited high expression of genes similar to disease-associated microglia (DAM)^31^, AMG4a and AMG1a subclusters most resembled DAMs (**Figure 3L; Table S3**). AMG4a exhibited particularly high expression of *Ank*, *Csf1*, and *Igf1* (**Figures 3M and 3N**), activation genes associated with Trem-2 dependent late-stage DAMs thought to have protective roles^31^. Like all other AMGs, AMG4a expressed high levels of injury-associated (**Figure 3C**) and lysosome-related genes (**Figure 3E**), but also exhibited uniquely high expression of phagocytic genes *Lpl*, *Cd9*, and *Myo1e* (**Figures 3M**, **3N, and S3E; Table S1**). This suggests that AMG4a may also be involved in phagocytosis despite myelin debris clearance already declining by 10dpl^59^. Moreover, we found that compared to AMG1a, AMG4a DEGs were enriched in activated pathways including B cell communication, and synapse pruning (**Figure 3O**). Taken together these data suggest that late-stage AMG4a microglia may be involved in adaptive immune crosstalk and synaptic reorganization, perhaps through the upregulation of a number of DAM-related genes (**Figures 3M**, **3N, and S3E**) such as *Cadm1* (**Figure S3P**), a cell adhesion molecule associated with synapse formation^60^ and glial interactions^61^.

AMG2 was most prominent at 10dpl and expressed both AMG1a/b and injury-associated BAM (iBAM) gene markers, including *Gpnmb* and *Atp6v0d2* (AMG1) and *Maf*, *Zbtb16*, and *Ms4a4a* (iBAM), suggesting a transitional activation state (**Figures 3A**, **3B and S3E; Table S1**). All BAM subclusters were defined by *Mrc1*, *Cd163*, and *F13a1* (**Figure S3E; Table S1**). iBAMs were distinguished from BAMs by the additional upregulation of genes associated with antigen-presentation (*H2-Aa*, *H2-Ab1*, and *Cd74*). Furthermore, the frequency of iBAMS in lesions increased as remyelination progressed, suggesting a potential role in antigen presentation to T cells during the late stages of remyelination (**Figures 3A**, **3B, and S3E**). AMG3 expressed non-unique marker genes (**Figure S3E**), also suggesting a transitional activation state. Additionally, AMG3 is largely absent by 10dpl (**Figures 3A and 3B; Table S2**), suggesting AMG3 may be early stage AMG4 microglia as they are highly correlated (**Figures S3F, S3G, and S3H**). Like AMG4a/b and iHMG, AMG3 expressed high levels of homeostatic genes found in HMGs (**Figures 3D and S3E**), suggesting these subclusters may represent steps towards a homeostatic state. These observations suggest that a shift in microglial activity towards a homeostatic program is initiated by 10dpl and increases as remyelination progresses.

### Astrocytes balance glial scar formation and angiogenesis throughout remyelination

Following LPC-mediated demyelination, astrocyte nuclei were reduced but recovered by 20dpl (**Figures 1C and S4A; Table S2**). As astrocytes are generally spared from LPC-injection and do not typically show large reductions in cell numbers^62^, the nuclei reductions we observed may be the result of other significant increases in cell populations, like immune cells (**Figure S3A; Table S2**). Strikingly evident, however, was the large transcriptional alterations observed in astrocytes throughout remyelination (**Figures 1E, S4B, and S4C**). Unsurprisingly, astrocytes were highly activated at 5dpl (**Figure S4C, center**), upregulating conventional activation markers such as *Gfap*. At 20dpl astrocytes remain highly activated (**Figure S4B and S4C**) but exhibit transcriptional differences from early (5dpl) remyelination (**Figure S4C, right**), suggesting that astrocytes in lesions take on differential over the time course of remyelination.

To further characterize the observed transcriptional alterations, astrocytes were subclustered and annotated based on previously reported markers^63^ and calculated marker genes (**Figures 4A**, **4B, and S4D; Table S1**). Two main groups of astrocyte transcriptional states were observed. Astro1 expressed *A2m*, *Cd44*, *Adamtsl1*, and *Slc8a1* (**Figures S4E, S4F, and S4G)** converging with markers for white matter fibrous astrocytes^63^. Astro2 expressed *Gpc5*, *Slc7a10*, *Slc6a11*, and *Gria2* (**Figure S4E, S4F, and S4G**), indicating a transcriptionally distinct astrocyte subcluster^63,64^. Naïve samples consisted almost exclusively of Astro1 and Astro2 subpopulations, with a higher level of Astro2 (**Figures 4A and 4B; Table S2**). By 5dpl, astrocytes developed an injury-associated phenotype (iAstro), denoted by the upregulation of several genes including stress response *Fkbp5*, cell growth-related *Azin1*, heterodimeric L-amino acid transporters *Slc3a2* and *Slc7a5*, inhibitor of apoptosis *Bcl2l1*, focal adhesion *Tns1*, ECM-related *Itih5*, angiogenesis-related GTPase *Rhoj*, synaptic vesicle membrane *Syt2*, and metal ion transporter *Slc39a14* (**Figures 4C**), that is maintained throughout all remyelination timepoints (**Figures 4A and 4B; Table S2**).

**Figure 4.**
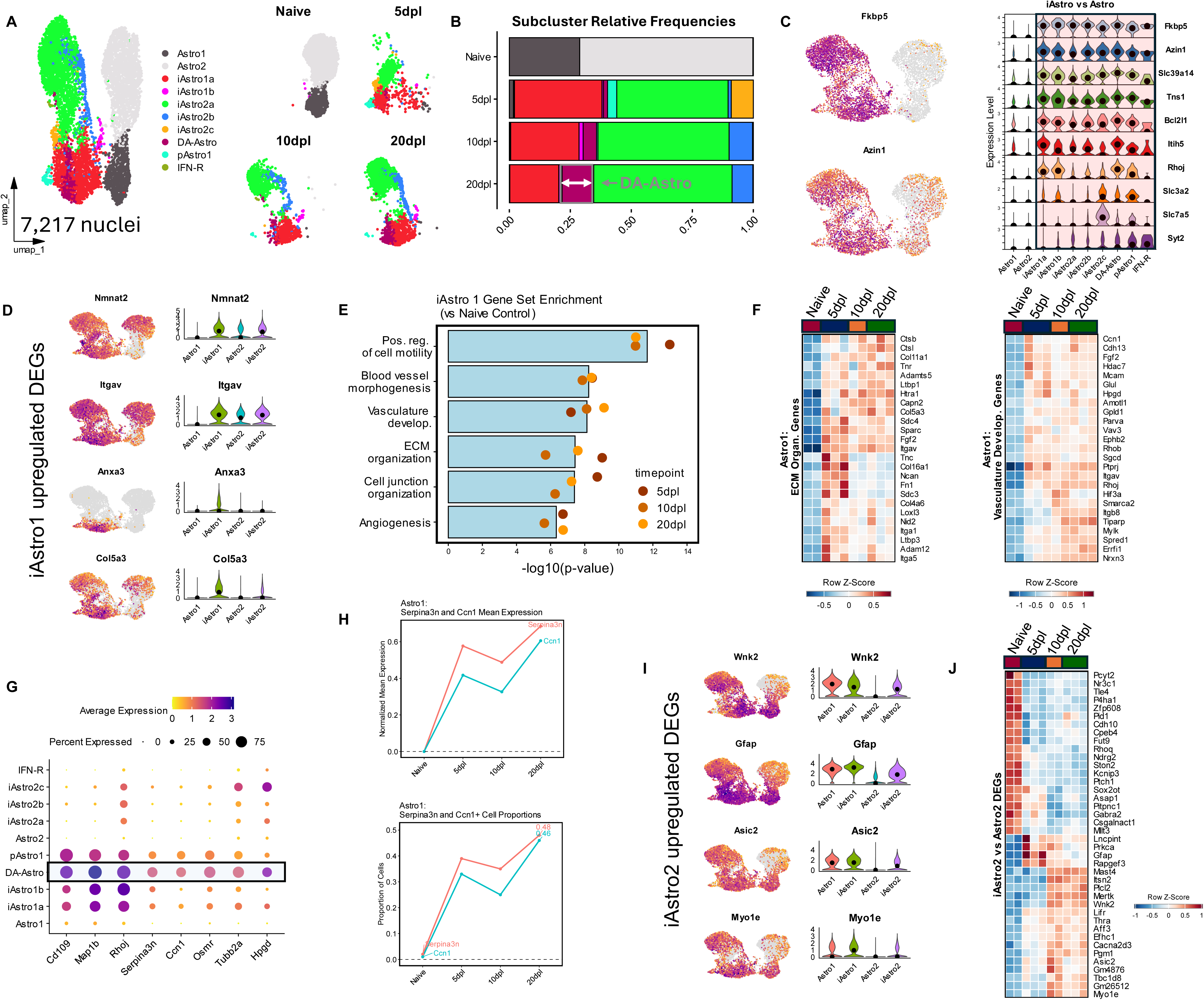
Injury-associated astrocytes are highly activated throughout remyelination, facilitating processes that may aid in remyelination progression. **A)** UMAP plot of 7,217 annotated astrocyte nuclei (*left*), split by timepoint (*right*). **B)** Bar plot showing the relative frequencies of annotated astrocyte subpopulations in each experimental timepoint, highlighting an increase in DA-Astro at 20dpl. See Table S2 for the number of nuclei and relative frequencies of all annotated clusters. **C)** Feature plot (*left*) and stacked violin plot (*right*) showcasing the expression of several pan injury-associated upregulated genes, including *Fkbp5* and *Azin1*, across astrocyte subclusters. Central dot in stacked violin plot represents the median expression. **D)** Feature plot (*left*) and violin plot (*right*) showcasing the expression of *Nmnat2*, *Itgav, Anxa3*, and *Col5a3*, select iAstro1 upregulated DEGs (compared to Astro1). Violin plots show the grouped expression of Astro1, iAstro1, Astro2, and iAstro2 subpopulations. Black dot in violin plot represents the median expression. **E)** Bar plot showcasing gene set enrichment terms associated with calculated upregulated DEGs between Astro1 astrocytes at each remyelination timepoint (5, 10, and 20dpl) vs naïve controls, organized by −log10(p-value). Bars represent the mean average of all three timepoints. Individual points represent the −log10(p-value) for 5dpl (*brown*), 10dpl (*light brown*), and 20dpl (*gold*) in each enrichment term. Gene set enrichment was conducted using Metascape. **F)** Scaled heatmaps showing differential gene expression patterns in Astro1 (Astro1, iAstro1a, iAstro1b, DA-Astro, pAstro1) nuclei across all experimental samples. Selected genes were acquired from gene sets for ECM organization (*left*) and vasculature development (*right*). **G)** Dot plot of all astrocyte subclusters highlight several activation genes upregulated in injury associated astrocytes with peak expression in DA-Astro, calculated form FindAllMarkers() in Astro1 subclusters. **H)** Trendlines indicating the normalized mean gene expression in (*top*) and relative proportion of (*bottom*) *Serpina3n* (*red*) and *Ccn1* (*blue*) Astro1 astrocytes across all experimental timepoints, indicating peak levels at 20dpl. **I)** Feature plot (*left*) and violin plot (*right*) showcasing the expression of *Wnk2*, *Gfap, Asic2*, and *Myo1e*, select iAstro2 upregulated DEGs (compared to Astro2). Violin plots show the grouped expression of Astro1, iAstro1, Astro2, and iAstro2 subpopulations. Black dot in violin plot represents the median expression. **J)** Scaled heatmap showing differential gene expression patterns in Astro2 (Astro2, iAstro2a, iAstro2b, iAstro2c, IFN-R) across all experimental samples. Selected upregulated genes were calculated from pairwise comparisons between all iAstro2 subclusters (iAstro2a, iAstro2b, iAstro2c, IFN-R) and naïve Astro2. Genes were further selected by top ranking based on adjusted p value.

Demyelination further led to the emergence of several subclusters at 5dpl, including proliferative astrocytes (pAstro1), disease-associated astrocytes (DA-Astros), and several iAstro subpopulations (**Figures 4A and 4B; S4D**). iAstro1 subclusters (iAstro1a, iAstro1b, pAstro1, DA-Astro) upregulated several genes including *Nmnat*, *Itgav*, *Anxa3*, and *Col5a3* (**Figure 4D**) Gene set enrichment analysis across all three remyelination timepoints suggests iAstro1 is highly mobile and involved in ECM remodeling and revascularization of the lesioned area (**Figures 4E and 4F**). ECM organization (**Figure 4E**) and several structural ECM genes including *Ncan*, *Fn1*, and *Tnc* (**Figure 4F, left**) peak at 5dpl, suggesting an early significant involvement in ECM deposition and glial scar formation^65,66^, where proteases such as *Ctsb* and *Ctsl* continue to be highly expressed at 20dpl (**Figure 4F, lef**t), suggesting continual maintenance of the glial scar^67^. Additionally, at 20dpl, there is an increased expression of lipid related *Apoe* and *St6galnac3* and glutamate signaling genes *Astn2* and *Ccser1* (**Figure S4H**), indicative of a potential return to homeostatic lipid functionality in iAstro1, supporting the later stages of remyelination. Interestingly, DA-Astros, demarcated by high expression of *Serpina3n* and *Ccn1* (**Figure 4G; Table S1**), were first observed at 5dpl but significantly increased as remyelination progressed (**Figures 4A and 4B**), much like DA-MOLb (**Figures 2A and 2B; Table S2**). DA-Astro is highly correlated with iAstro1a (**Figure S4I**), shares core activation genes with other iAstro1 subclusters (**Figures 4G and S4D**) and appears to be strongly involved in ECM remodeling and revascularization (**Figure S4J**). While it is not yet known whether DA-Astros are beneficial to or detrimental to remyelination, DA-Astro predicted functionality and the fact that *Serpina3n* and *Ccn1* mean expression and cell positive proportions peak at 20dpl (**Figure 4H**), suggests DA-Astros may support remyelination.

On the other hand, iAstro2 subclusters (iAstro2a, iAstro2b, iAstro2C, IFN-R) selectively upregulated genes such as *Wnk2*, associated with cell migration, *Gfap*, intermediate filament protein associated with astrogliosis, *Asic2*, a sodium channel component and *Myo1e*, related to endocytosis (**Figure 4I**). Interestingly, while these genes were highly expressed in homeostatic Astro1 and iAstro1 they are either absent or expressed at low levels in homeostatic Astro2 (**Figure 4I**), indicating reactive heterogeneity. This finding is particularly notable since *Gfap* is often used as a general marker of astrogliosis^68^, yet our results indicate it specifically marks reactivity in iAstro2 subclusters. Additionally, iAstro2 populations, while highly activated, upregulated fewer unique DEGs compared to iAstro1 (**Figure S4K**). Moreover, iAstro2 exhibited differences in DEGs between 5dpl and later remyelination stages at 10 and 20dpl (**Figure 4J**). This suggests differential roles during remyelination with *Gfap* marking activity at 5dpl, and genes like *Mertk*, *Wnk2*, and *Myo1e* at later stages, indicating potential phagocytic roles astrocytes associated with late stage remyelination.

### Fibrotic scar formation and vascular reprogramming observed throughout remyelination

We found that LPC-mediated focal demyelination also had profound effects on vascular and mesenchymal cell populations in the spinal cord. While total nuclei numbers were similar across experimental conditions (**Figure S5A**), transcriptional alterations, most notably at 5dpl, were observed in lesion samples (**Figures S5B and S5C**). These data suggest that early following demyelination, vascular and mesenchymal cells are highly activated, and that over time those reactive profiles alter as remyelination progresses (**Figure S5C**). To further analyze transcriptional alterations, we subclustered and annotated vascular and mesenchymal nuclei using both previously reported markers^17^ and calculated cluster marker genes (**Figures 5A and S5D; Table S1**). Subclustering revealed several fibroblast subpopulations (Fibro.1-4), endothelial cells (Endo. 1a, 1b and 2), pericytes, ependymal cells, and smooth muscle cells (SMCs) (**Figure 5A**). We observed a striking increase in meningeal fibroblasts (Fibro. 1 clusters) at 5dpl that reduced over time but remained higher than naïve condition at 20dpl (**Figures 5A and 5B; Table S2**). Inducing LPC-lesions requires penetrating the meninges to deliver the injection into the mouse spinal cord^18^. This suggests that meningeal fibroblasts invade the lesion site^69^ and begin to accumulate early after injury. By contrast, perivascular fibroblasts (Fibro. 2-4 clusters), pericytes, and ependymal cells in lesions were lower at 5dpl compared to naïve samples but returned to near homeostatic frequencies by 20dpl (**Figures 5A and 5B; Table S2**), indicating a recovery from blood brain barrier (BBB) disruption. Fibroblasts at 5dpl were further subclustered into early activated Fibro. 1 (EA-Fibro. 1), proliferative EA-Fibro.1 (pEA-Fibro. 1), and EA-Fibro. 4 (**Figures 5A and 5B; Table S2**). These subclusters were enriched in fibrosis-related *Runx1*, *Runx2*, several collagen genes, and matrix metalloproteinases (MMPs) *Adam19* and *Bmp1* (**Figure 5C**). Collagen is an abundant component of the fibrotic scar, which forms in MS, EAE, and LPC-mediated demyelination^70–73^. While several collagen genes and extracellular matrix (ECM) proteins were upregulated throughout remyelination (**Figure 5D**), we found that they were most highly enriched in fibroblasts at 5dpl (**Figures 5D and 5E**), suggesting that fibrotic scar formation primarily occurs at 5dpl following demyelinating injury, facilitated by both meningeal and perivascular fibroblasts (**Figures 5C and S5E**). Fibroblasts were dynamic and exhibited significant differential gene expression between experimental timepoints (**Figure S5F**). In meningeal and perivascular fibroblasts, many DEGs that define early (5dpl) and late (20dpl) remyelination were shared (**Figure 5F**), further suggesting similar time-dependent functions in both groups. Indeed, at 10 and 20dpl, several genes related to cholesterol processing and glucocorticoid (GC) and corticosteroid (CS) signaling are upregulated across fibroblasts (**Figure S5G**), perhaps directly aiding in remyelination as newly generated OLCs rely on cholesterol processing^14^. In meningeal fibroblasts, upregulated genes such as *Slc47a1*, *Slc47a2*, and *Slc23a2* suggest a return to homeostatic functionality while *Thsd7a* and *Glul*, suggest an increase in angiogenesis at 10 and 20dpl (**Figure 5G**). In perivascular fibroblasts, activation genes such as *Bbs9*, *Slc38a2* and *Arhgap18* are prominent at 20dpl (**Figure 5H**), suggesting these cells are still highly reactive.

**Figure 5.**
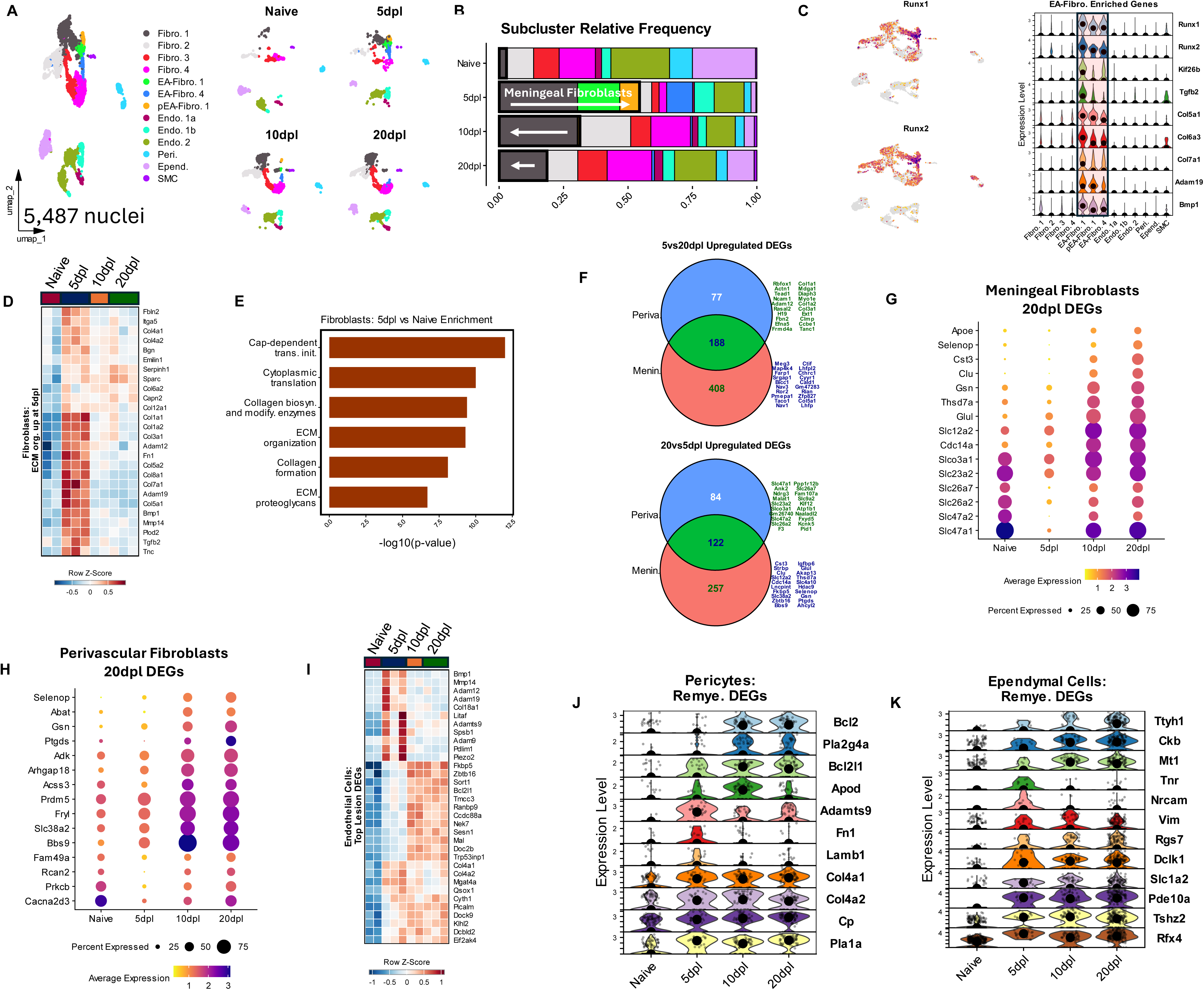
Fibrotic scar formation and vascular/mesenchymal reprogramming evident throughout remyelination. **A)** UMAP plot of 5,487 annotated vascular and mesenchymal nuclei (*left*), split by timepoint (*right*). **B)** Bar plot showing the relative frequencies of annotated vascular/mesenchymal subpopulations in each experimental timepoint, highlighting the expansion of meningeal fibroblasts (Fibro. 1, EA-Fibro. 1, and pEA-Fibro. 1) in lesions. See Table S2 for the number of nuclei and relative frequencies of all annotated clusters. **C)** Feature plot (*left*) and stacked violin plot (*right*) showcasing the expression of EA-Fibro. enriched genes including several fibrosis-related genes including *Runx1* and *Runx2*, multiple collagen genes, and matrix metalloproteinases *Adam19* and *Bmp1* across all vascular/mesenchymal subclusters. Central dot in stacked violin plot represents the median expression. **D)** Scaled heatmap showing differential gene expression patterns in all fibroblasts across experimental samples. Selected genes were acquired from gene sets for ECM organization enriched at 5dpl. **E)** Bar plot showcasing gene set enrichment terms associated with calculated upregulated 5dpl DEGs from comparison of fibroblasts at 5dpl vs naïve controls, organized by −log10(p-value). Bars represent the −log10(p-value) for each enrichment term. Gene set enrichment was conducted using Metascape. **F)** Venn diagrams showcasing the overlap in upregulated DEGs in meningeal and perivascular fibroblasts at 5dpl (*top*) and 20dpl (*bottom*) calculated from comparing 5 vs 20dpl in meningeal and perivascular fibroblasts, respectively. The top 20 upregulated genes unique to meningeal fibroblasts at 5 and 20dpl (*top green text*) and shared between both fibroblast groups (*bottom blue text*) are shown. **G)** Dot plot of meningeal fibroblasts (Fibro. 1, EA-Fibro. 1, and pEA-Fibro. 1) grouped by experimental timepoint highlighting several genes upregulated at 20dpl, calculated from 20dpl vs 5dpl DEG testing. **H)** Dot plot of perivascular fibroblasts (Fibro. 2, Fibro. 3, Fibro. 4, and EA-Fibro. 4) grouped by experimental timepoint highlighting several genes upregulated at 20dpl, calculated from 20dpl vs 5dpl DEG testing. **I)** Scaled heatmap showing top differential gene expression patterns in all endothelial cells across experimental samples. Selected genes were acquired from DEG tests comparing all lesion and naïve timepoints to each other to highlight activation differences at 5 and 20dpl. **J)** Stacked violin plot showcasing the expression of top remyelination genes (*lesion timepoints compared to naïve controls*) enriched in pericytes. Central dot in stacked violin plot represents the median expression. **K)** Stacked violin plot showcasing the expression of top remyelination genes (*lesion timepoints compared to naïve controls)* enriched in ependymal cells. Central dot in stacked violin plot represents the median expression.

We found that endothelial cells (Endo. 1a, Endo.1b, and Endo. 2) exhibited fewer relative frequency changes (**Figure 5B; Table S2**) and DEGs upon lesioning (**Figure S5H**). We observed at 5dpl an upregulation of several activation-related genes such as collagen genes *Col4a1*, *Col4a2*, and *Col18a1* as well as several MMPs including *Adam9*, *Adam12*, *Mmp14*, *Bmp1*, *Adam19*, and *Adamts9* (**Figure 5I**), indicating early involvement in ECM and vascular remodeling. By later timepoints 10 and 20dpl, we observed increases in genes such as BBB integrity and clearance protein *Picalm*, anti-apoptotic *Bcl2l1*, sphingolipid/ceramide-related *Sort1*, Akt signaling activator *Tmcc3*, and stress response *Trp53inp1* and *Sesn1* (**Figure 5I**), suggesting that endothelial cells continue to remodel and possibly recover lost vasculature as well as initiate pro-survival functionality. Despite these differences, endothelial cells at early and late stages of remyelination are fairly similar (**Figure S5H**), suggesting that they maintain a similarly activated state throughout remyelination. Moreover, similar to other vascular and mesenchymal cell populations, pericytes exhibited high expression of activation (*Fn1* and *Lamb1*) and ECM remodeling (*Adamts9*, *Col4a1*, and *Col4a2*) genes at 5dpl, while upregulating pro-survival gene expression (*Bcl2*, *Bcl2l1*) by 10dpl. Interesting, pericytes upregulated phospholipases *Pla1a* and *Pla2g4a*, the ferroxidase *Cp*, and the secreted glycoprotein *Apod*, which is known to be important for remyelinating OLs^74^, suggesting that pericytes may be interacting with myelin debris or OLCs directly (**Figure 5J**). Additionally, we found that ependymal cells upregulated and maintained high expression of cilia-related *Rfx4*, cell growth regulator *Tshz2*, phosphodiesterase *Pde10a*, excitatory amino acid transporter *Slc1a2*, pro-survival and Wnt regulator *Dclk1*, GAP protein *Rgs7*, and structural activation protein *Vim* throughout remyelination (**Figure 5K**). Cell adhesion molecules *Nrcam* and *Tnr* were most highly expressed at 5dpl suggesting an early involvement in establishing structural support. Metallothionein *Mt1*, creatine kinase *Ckb*, and chloride channel *Ttyh1* were most highly expressed at later remyelination timepoints, suggesting increasing involvement in metal ion metabolism and creatine production, both of which could support newly generated, remyelinating oligodendrocytes^75,76^.

Together these data indicate several vascular and mesenchymal cell subpopulations may directly support remyelination, potentially by initial fibrosis of the demyelinated area by 5dpl (**Figure S5E**), led by meningeal and perivascular fibroblasts (**Figures 5C and 5D**), and increased activation (**Figure S5I**). The extent to which angiogenesis, metal ion transport, and cholesterol/lipid processing influence late-stage remyelination remains to be determined. However, by 20dpl, multiple cell populations seem to activate pro-survival mechanisms (**Figures S5J**), which may be critical for the successful completion of remyelination.

### Nonlesion control samples reveal systemic generalized activation caused by a single focal demyelinating event

In addition to lesioned and naïve spinal cord white matter tissue, we also collected adjacent nonlesioned spinal cord tissue as internal controls for comparison and analyzed the data in parallel (**Figures 1A and S6**). We found that the nonlesioned samples displayed similar numbers of nuclei, similar QC metrics, and no obvious technical batch effects at all three remyelination timepoints analyzed (**Figure S6A-S6C**). Moreover, nonlesioned samples, when compared to each other, exhibited few DEGs at all remyelination timepoints (**Figure. S6D**). With the exception of Schwann cells, all major cell populations found in naïve and lesioned samples were also observed in nonlesioned samples (**Figures 1B and S6E**), and with similar relative proportions (**Figure S6F; Table S2**). We found that nonlesioned samples exhibited a large number of DEGs compared to naïve samples (**Figure S6G**). Compared to lesioned samples, however, nonlesioned samples only exhibited significant DEGs in immune and vascular nuclei (**Figure S6H**). These data indicate that nonlesioned samples differed transcriptionally from naïve controls, suggesting cellular populations outside of directly lesioned areas are systematically activated by focal demyelinating injury. However, strong vascular and immune activation patterns appear to be unique to remyelinating lesion sites.

We next subclustered OLC, immune, astrocyte, and vascular nuclei and analyzed transcriptomic alterations in the nonlesioned samples. In OLCs, we found that all mature OL nuclei were injury-associated (**Figures S6I and S6J**), matching the lesioned samples (**Figures 2A and 2B**). The injury-associated state involves the upregulation of several activation genes seen across all nonlesion and lesion samples (**Figure S6K**), coupled with the downregulation of several myelin related genes found most expressed in naïve MOLs (**Figure S6L**). The injury-associated activated state was observed in nonlesioned samples with no indication of an upregulation in OLC differentiation (**Figures S6J**), suggesting that the consequences of the shared activation pattern in lesioned and nonlesioned tissues differ. In immune cells, we found no evidence of AMG1a/b macrophage-like subclusters (**Figure S6M; Table S2**). AMG1a/b nuclei are characteristic of early timepoint 5dpl in lesions (**Figures 3A and 3B**) and are responsible for high levels of classic inflammatory activity (**Figure 3J**) marked by phagocytosis-related *Atp6v0d2* and *Gpnmb* (**Figures 3I**), both of which are not expressed in nonlesioned samples (**Figure S6N**), indicating that the early accumulation of macrophage-like microglia is unique to the lesion site. Of note, nonlesioned microglia are slightly activated compared to naïve nuclei (**Figure S6N**) but also exhibited high expression of several homeostatic genes. Similarly to OLCs, nonlesioned astrocytes acquired an injury-associated phenotype at all remyelination timepoints that resembled lesioned astrocytes (**Figures S6O and S4C**). Utilizing *Serpina3n* and *Ccn1* as markers of highly activated Astro1 populations, we found that nonlesioned Astro1 upregulated both genes in a similar pattern to lesioned Astro 1, but to only half the extent (**Figures S6P, S6Q, and 4H**). These data suggest that astrocytes are activated systemically due to demyelinating injury, but that systemic activation is not as strong away from the lesion stie. Notably, among vascular and mesenchymal cells in nonlesioned samples, no evidence for increased meningeal fibroblasts was observed (**Figure S6R and S6S**). Since meningeal Fibro. 1 exhibit upregulated expression of fibrotic scar/fibrosis-related genes (**Figure 5C**), their lack of detection in nonlesioned samples indicates fibrotic scar formation is also unique to the lesion site. Further, endothelial cells (**Figure S6T**), pericytes (**Figure S6U**), and ependymal cells (**Figure S6V**) in nonlesioned samples exhibited similar, albeit weaker, activation profiles as lesioned samples. Collectively, these findings indicate that a single focal demyelinating injury induced by LPC activates all major glial cell types in nonlesioned tissues beyond the directly injured site. This suggests that glial cells react to systemic inflammation, which can occur distally to the injury site and persist for weeks.

## DISCUSSION

The LPC-induced focal demyelination mouse model is extensively used to study the process of remyelination and to test various pharmacological and genetic interventions for their effect on remyelination efficacy^12,18,39^. While this model lacks all adaptive immune responses characteristic of MS, it recapitulates many microenvironmental elements including T lymphocyte recruitment, reactive gliosis, glial and fibrotic scarring, and axonal dysregulation^12,18,39^. Importantly, the LPC focal demyelination model is a system in which efficient and successful remyelination is inherent. Therein, the various immune components, glial activation states, and overall expression shifts following demyelination in this model are all expected to support acute, spontaneous remyelination within a defined timeframe. Furthermore, the LPC model enables assessments of cellular changes at highly reproducible time points to determine when and how remyelination is facilitated. Transcriptomic analyses of LPC-induced remyelination at the single cell and single nucleus level have been conducted, focused primarily on microglia^13^ or oligodendrocytes^35^ in isolation. Here, we present a holistic transcriptomic analysis of multiple cell types throughout remyelination in the LPC-mediated demyelination model, defining cell type-specific signatures and activation states in homeostatic tissue, during three stages of remyelination, and in adjacent nonlesioned tissue in order to better understand the cellular alterations necessary to facilitate remyelination in mice.

OLCs had a multifaceted response to LPC-induced demyelination. As early as 5dpl, we found evidence at the transcriptional level for OPC differentiation, which is earlier than typically reported by others^18,77,78^. Additionally, at 5dpl both immature OPCs we believe are found inside the lesion and perilesional MOLs acquired a disease-associated (DA) signature marked by *C4b*, *Serpina3n*, and *Syt4*. DA-OLCs have been previously observed in MS-related mouse models and across various CNS injuries and diseases^17,34,35,43^, but what role they play in the remyelination process is not yet well understood. Importantly, DA-OPCs and COPs are only found in lesioned samples at 5dpl, suggesting that these subpopulations are not only lesion-specific but perhaps also tied to early stages of inflammatory signaling^79,80^. Further, we found that all injury-associated MOLs from lesioned and nonlesioned tissue, which persist through 20dpl, exhibited an increase of several activation markers that appear to be highly expressed in immature OLCs, but that we found are downregulated in MOLs in homeostatic tissue. Concurrently, injury-associated MOLs also exhibited decreases in several myelin-related genes that were expressed at high levels in naïve MOLs, whereas more traditional myelin genes were not affected^81^. Taken together, these data suggest that MOLs not only become reactive following demyelination, but that MOL reactivity may interfere with the expression levels of several myelin-related genes. Teasing apart the role of these activation genes, be it to facilitate immune communication, to permit survival and maturation as suggested of other genes that are upregulated in newly generated MOLs during remyelination^14,76^, or as indicators of stress-induced alterations that interfere with myelin gene expression is yet to be determined. Overall, these phenomena may help explain why myelin often appears thinner in remyelinating lesions and may provide potential targets for pharmaceutical intervention.

Following LPC-mediated demyelination, microglia migrate and proliferate at the lesion site, and upregulate multiple genes involved in phagocytosis and lipid storage and processing, suggesting that throughout remyelination microglial activation involves some form of lysosomal processing. At early timepoint 5dpl, accumulation of microglia was prominent and was dominated by the presence of macrophage-like, Lys2+ microglia. We found that these cells were highly enriched in genes regulating intracellular pH reduction, neutrophil degranulation, and autophagy, suggesting they are classical pro-inflammatory microglia. Macrophage-like microglia are vastly reduced by 10dpl, suggesting they may be involved in early myelin debris removal, after which they are displaced. It has been suggested this may be an inevitable characteristic of remyelination where necroptosis of an early responsive microglia population enables replacement by pro-regenerative microglia^29^. Contextually, this is important as it provides a backdrop for our findings at later stages of remyelination, notably 20dpl, where microglia are observed to be in an activated state modulating components of the adaptive immune system and also upregulating several homeostatic genes, suggesting they may be returning to homeostasis and at the same time may be providing crucial pro-regenerative roles to support remyelination.

Like OLCs, astrocytes responded to LPC-mediated demyelination both at the lesion site and in adjacent, nonlesioned tissue. Previously described in other studies^34^, astrocytes developed a disease(demyelination)-associated signature reminiscent of DA-OLCs, demarcated by the expression of *C4b* and *Serpina3n*. Notably, the disease-associated phenotype in astrocytes continued through and peaked at 20dpl both in lesioned and nonlesioned tissues. Reactive astrocyte populations were heavily involved in ECM organization and angiogenesis, suggesting that after LPC-mediated focal demyelination, astrocytes contribute to glial scar formation and re-vascularization of the lesion site. Our data also highlighted robust alterations in vascular and mesenchymal cells in response to demyelination and throughout remyelination. Fibroblasts, especially meningeal fibroblasts, dramatically increase in the lesion following focal demyelination and remain at high numbers throughout remyelination. DEG testing and gene set enrichment analyses suggest that fibrotic scar formation and maintenance were prominent features of fibroblasts found early in the remyelination process. While fibroblast accumulation remained high, collagen-related genes and pathways were downregulated by 10 and 20dpl, while cholesterol related genes were upregulated. Cholesterol processing is crucial to newly generated MOLs during remyelination^14^, therein suggesting that fibroblasts may support the generation of new MOLs directly. The unique characteristics we observed at the lesion site, including the expansion of macrophage-like microglia and meningeal fibroblasts, suggest that a specific combination of various cell populations defines the environment in which remyelinating oligodendrocytes are generated. These factors are critical to consider in the context of the remyelination process

As previously described, our results also indicate that there are transcriptomic alterations induced in adjacent, nonlesioned white matter tissue resulting from LPC-mediated focal demyelination. in this study. However, utilizing high throughput single nucleus RNA sequencing, we show in the current study strong evidence for a generalized demyelination-associated injury signature in both lesioned samples and nonlesioned controls. In MS, normal appearing white matter (NAWM) also displays nontypical characteristics similar to lesioned areas^82–84^, suggesting that in MS systemic inflammation also influences surrounding tissues. Further research is needed to determine whether systemic activation of glial cells has harmful effects distant from the demyelinating injury site.

Overall, our study provides a useful transcriptomic dataset for comparing cell type signatures in homeostatic tissue and throughout successful remyelination following LPC-mediated demyelination. We identified possible mechanisms by which activated responses from multiple cell types including OLCs, microglia, astrocytes, and fibroblasts facilitate a spontaneous remyelination in mice following focal demyelination, enriching our understanding of microenvironmental alterations underlying the remyelination process. Altogether, this study provides a valuable resource enabling the investigation of the holistic underpinnings of remyelination progression.

## Acknowledgements

This study was supported by the National Institutes of Health (2R56NS107523-06, 5R01NS107523), National Multiple Sclerosis Society Harry Weaver Neuroscience Scholar Award (JF-1806-31381), and Congressionally Directed Medical Research Programs (W81XWH-17-1-0268) to J.K.H., and the Patrick Healy Graduate Fellowship to G.S.M. We thank all members of our lab (La FamiGlia) for their intellectual comments and contributions to the analysis of the data for this manuscript.

## Author contributions

G.S.M, M. B., and J.K.H. designed the study. G.S.M. performed all mouse experiments and bioinformatic data analysis. Z.M., J.H., and M.B. contributed to lysolecithin lesions, nuclei isolation, and library preparation for snRNAseq. Z.M. and J.H. contributed to immunostaining. G.S.M. drafted the manuscript. G.S.M. and J.K.H. edited the manuscript. J.K.H. oversaw the study.

## SUPPLEMENTAL FIGURE LEGENDS

**Figure S1.**
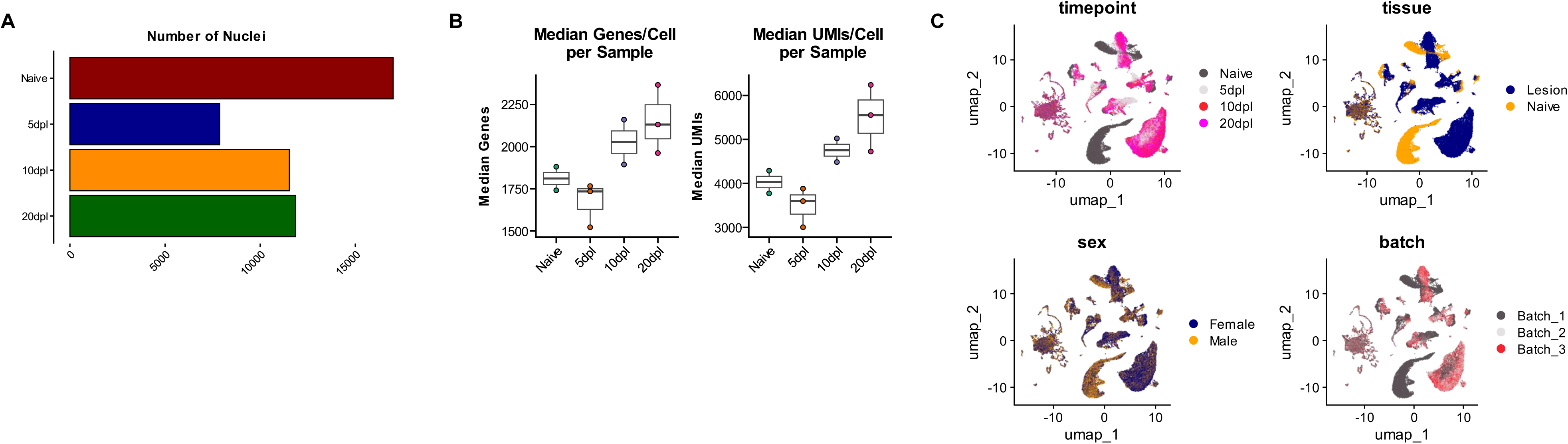
Characterization and quality control assessment of overview data. **A)** Bar plot showing the number of overall nuclei per timepoint, highlighting reductions in lesion timepoints. **B)** Box and whisker plots showing the median genes/Cell per experimental sample (*left*) and median UMIs/Cell per experimental sample (*right*) across timepoint conditions: Naïve, 5dpl, 10dpl, and 20dpl. **C)** UMAP projections split (clustered) by timepoint (*top left*), tissue (*top right*), sex (*bottom left*) and batch (*bottom right*), indicating together the lack of technical batch effects.

**Figure S2.**
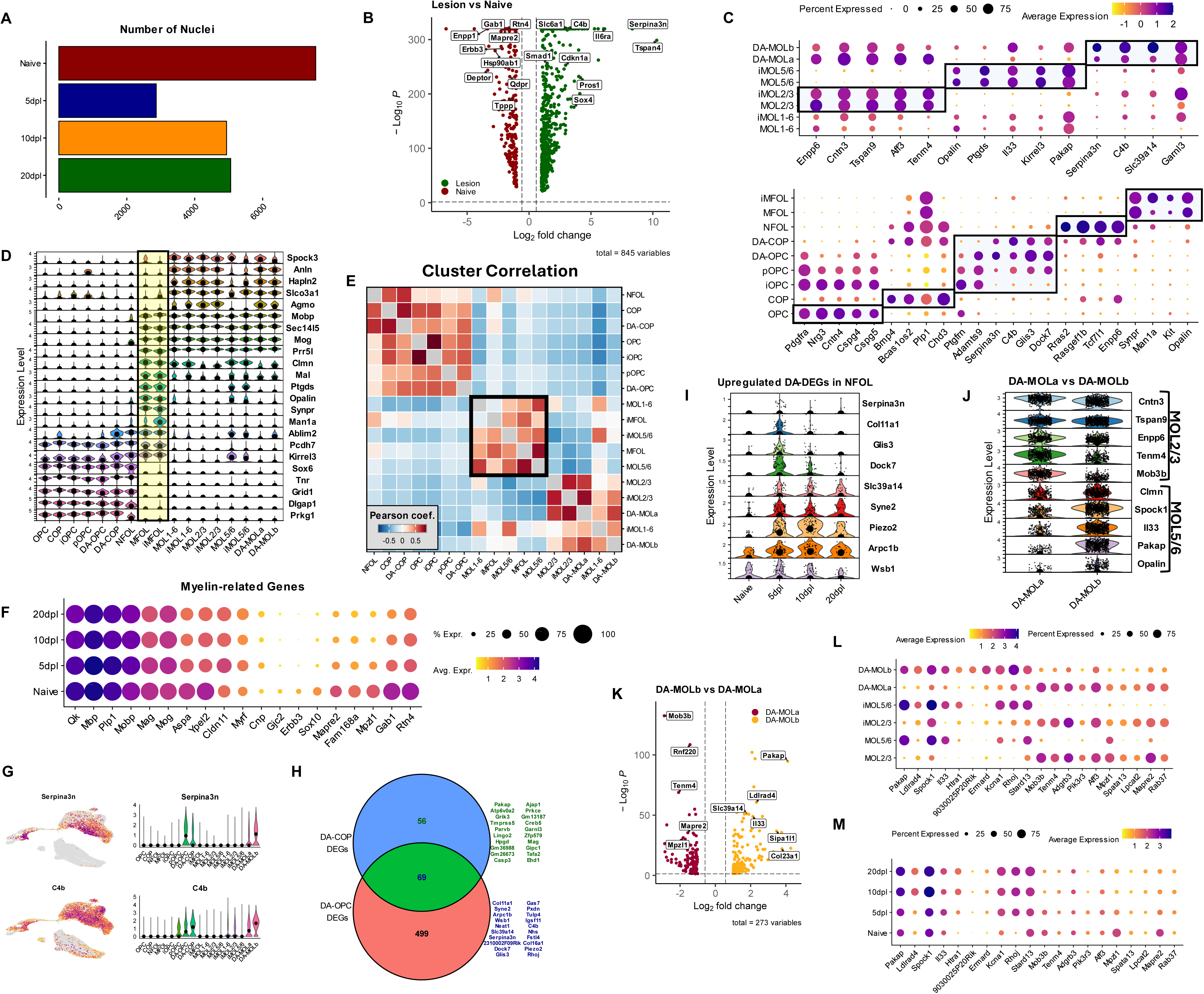
Characterization of oligodendrocyte lineage cells following LPC-mediated demyelination. **A)** Bar plot showing the number of OLC nuclei per timepoint, highlighting overall reductions in lesion timepoints. **B)** Volcano plots showcasing DEGs calculated from a pairwise comparisons of naïve OLCs (*red*) and lesion OLCs (*green*). Log2FC >1.0, min. pct = 0.2, adjusted p-value <0.05, MAST, Bonferroni correction. **C)** Dot plots showcasing top marker genes used to annotate both mature oligodendrocytes (*top*) and immature OLC populations (*including MFOLs, bottom*). See Table S1 for a complete list of cluster markers. **D)** Stacked violin plot showing several marker genes that distinguish MFOLs (*highlighted in the middle yellow box*) as a unique OLC subpopulation, expressing both immature, mature and unique genes. Central dot indicates the median expression of each gene. **E)** Heatmap showing the correlation (Pearson) between annotated subclusters based on the top 2000 variable genes. The black outline box highlights MFOLs and subclusters to which they are most correlated. **F)** Dot plot of mature OL subclusters showcasing the expression pattern of several-myelin related genes across naïve and lesion experimental timepoints. **G)** Feature plot (*left*) and violin plot (*right*) showcasing the expression of disease-associated genes *Serpina3n* and *C4b* amongst annotated OLC subclusters. Central dot in violin plot represents the median expression. **H)** Venn diagram showcasing the overlap (69 genes) in upregulated DEGs in DA subclusters calculated from pairwise comparisons of DA-COPs vs COPs (56 unique) and DA-OPCs vs OPCs (499 unique). The top 20 upregulated genes unique to DA-COPs (*top*) and shared between the DA groups (*bottom*) are shown. **I)** Stacked violin plot showing the upregulated expression of several DA-associated DEGs in lesion-specific NFOLs. Central dot shows the median gene expression. **J)** Stacked violin plot showing the enrichment of MOL2/3 and MOL5/6 marker genes in DA-MOL subclusters, indicating that DA-MOLa is predominantly MOL2/3, while DA-MOLb is mixed. Central dot represents median gene expression. **K)** Volcano plot showcasing DEGs calculated from a pairwise comparisons of DA-MOLb (*gold*) and DA-MOLa (*maroon*). Log2FC >1.0, min. pct = 0.2, adjusted p-value <0.05, MAST, Bonferroni correction. **L)** Dot plot of mature OL subclusters showcasing top DEGs calculated form pairwise comparisons between DA-MOLb and DA-MOLa. **M)** Dot plot of mature OL subclusters grouped by experimental timepoint showcasing top DEGs calculated form pairwise comparisons between DA-MOLb and DA-MOLa.

**Figure S3.**
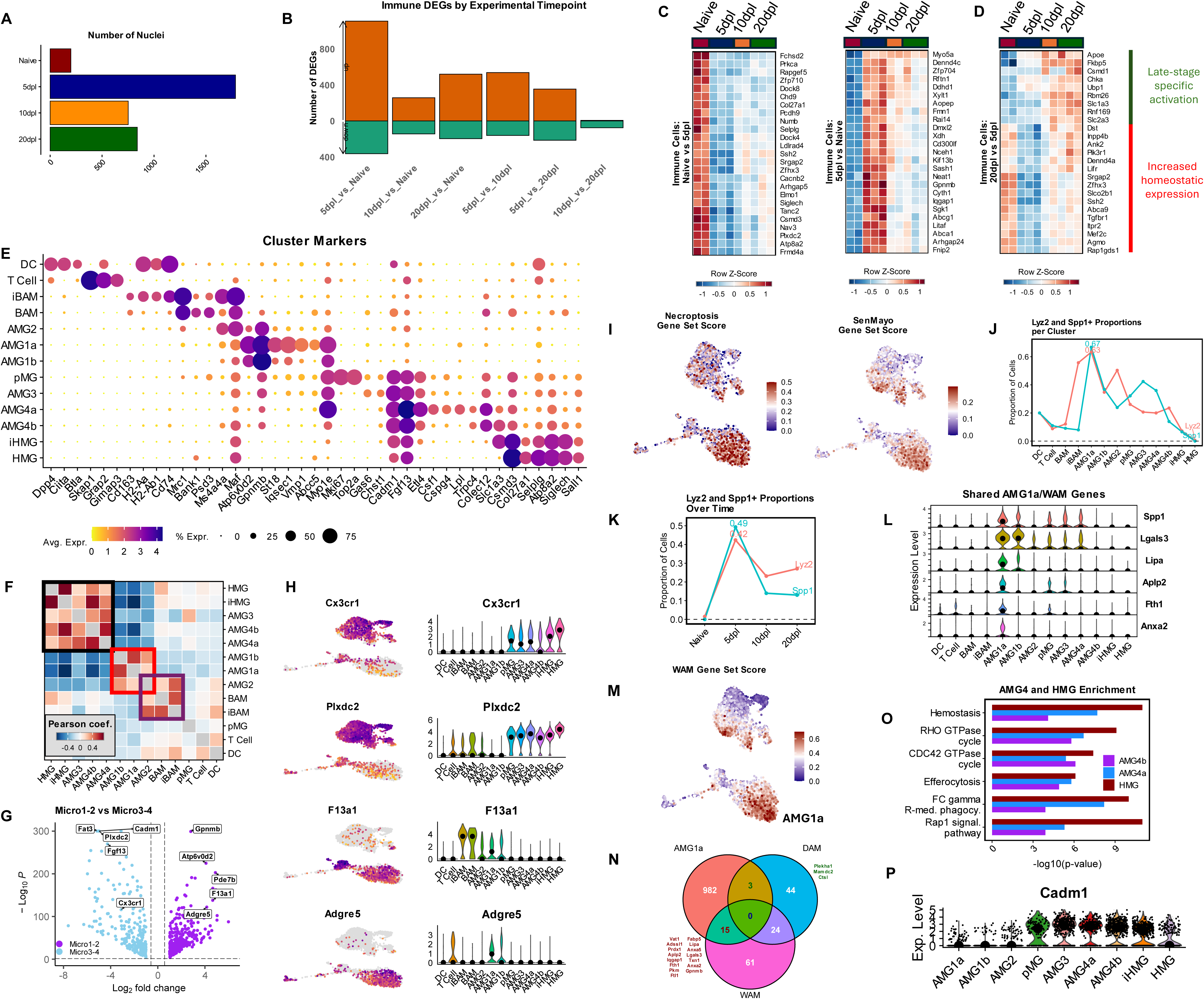
Characterization of immune cells in lysolecithin mouse model. **A)** Bar plot showing the number of immune nuclei per timepoint, highlighting large increases at lesion timepoints. **B)** Stacked bar plot indicating the number of upregulated (*top, orange*) and downregulated (*bottom, green*) genes calculated in pairwise comparisons in all immune cells between each experimental timepoint (5dpl, 10dpl, 20dpl, and Naïve). Log2FC >1.0, min. pct = 0.2, adjusted p-value <0.05, MAST, Bonferroni correction. **C-D)** Scaled heatmaps showing gene expression patterns in immune nuclei from all samples. Expression of upregulated naive DEGs, calculated from 5dpl vs Naïve (*C, left*), of upregulated 5dpl activated DEGs (*C, right*), and upregulated 20dpl DEGs, calculated from 5dpl vs 20dpl. (D) highlights concurrent alterations in activation and increase in homeostatic gene expression. Genes were selected by top ranking based on adjusted p value. **E)** Dot plot showcasing top marker genes used to annotate immune subclusters. See Table S1 for complete markers. **F)** Heatmap showing the correlation (Pearson) between annotated subclusters based on the top 2000 variable genes. The black outline box highlights microglial clustering (AMG3, AMG4, iHMG, HMG), the red box highlights macrophage-like microglial clustering (AMG1, AMG2), and the purple box highlights AMG2s association with both BAMs and macrophage-like clusters. **G)** Volcano plot showcasing DEGs calculated from a pairwise comparison of AMG1-2 (*purple*) vs AMG3-4 (*blue*), highlighting differences between macrophage-like microglia and microglial clusters. Log2FC >1.0, min. pct = 0.2, adjusted p-value <0.05, MAST, Bonferroni correction. **H)** Feature plot (*left*) and violin plot (*right*) showcasing the expression of AMG3-4 markers *Cx3cr1* and *Plxdc2* (*top*), and AMG1-2 markers *F13a1* and *Adgre5* (*bottom*). Central dot in violin plot represents the median expression. **I)** Feature plots showing necroptosis (*left*) and SenMayo (*right*) gene set scores projected onto the immune UMAP. Table S3 for the complete gene set lists used. **J)** Trendline showing the proportion of *Lyz2* and classical early DAM marker *Spp1* positive cells in annotated immune clusters. **K)** Trendline showing the proportion of *Lyz2* and classical early DAM marker *Spp1* positive across naïve and lesion timepoints, highlighting steep declines between 5 and 10dpl. **L)** Violin plot highlighting key WAM gene expression in immune annotated clusters. AMG1a exhibited the highest expression of several WAM genes. Central dot represents the median expression. **M)** Feature plot highlighting calculated WAM gene set scores on the projected immune UMAP. AMG1a exhibited the highest average levels of WAM gene set expression. See Table S3 for the complete gene set lists used. **N)** Venn diagram showing the overlap of AMG1a cluster markers with WAM and DAM gene sets. The genes shared between AMG1a and DAM (*green*) and AMG1a and WAM (*red*) are shown. FindAllMarkers() was used to determine cluster markers. See Table S3 for the complete gene set lists used. **O)** Bar plot showcasing gene set enrichment terms associated with calculated upregulated DEGs shared between AMG4 and naïve HMG vs highly activated AMG1a, organized by −log10(p-value). Bars represent the −log10(p-value) for upregulated DEGs in HMG (*darkred*), AMG4a(*blue*), and AMG4b(*purple*). Gene set enrichment was conducted using Metascape. **P)** Violin plot highlighting the expression of *Cadm1* amongst annotated subclusters. Central dot represents median expression.

**Figure S4.**
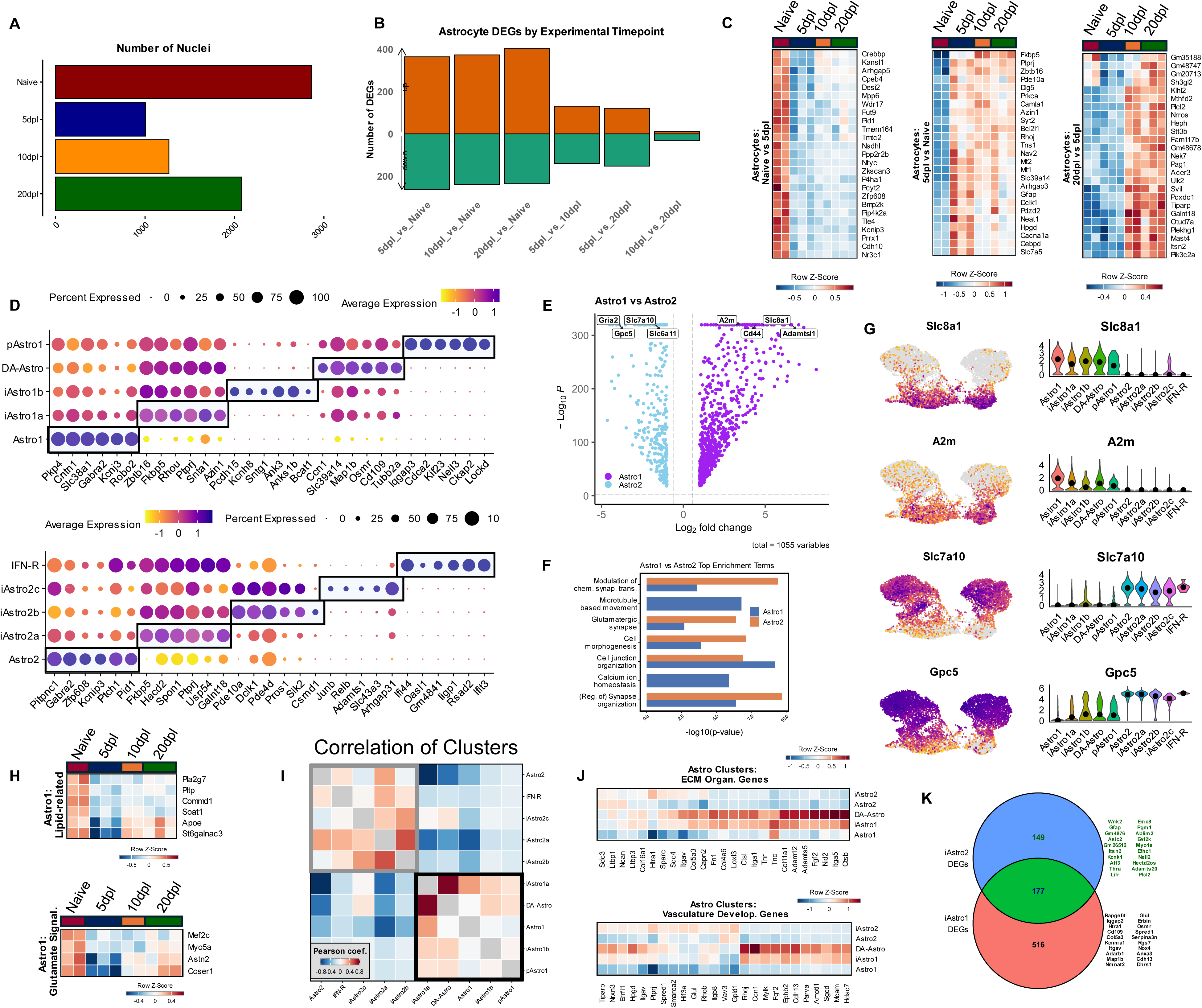
Characterization of astrocyte subclusters throughout remyelination. **A)** Bar plot showing the number of astrocyte nuclei per timepoint. **B)** Stacked bar plot indicating the number of upregulated (*top, orange*) and downregulated (*bottom, green*) genes calculated in pairwise comparisons in all astrocytes between each experimental timepoint (5dpl, 10dpl, 20dpl, and Naïve). Log2FC >1.0, min. pct = 0.2, adjusted p-value <0.05, MAST, Bonferroni correction. **C)** Scaled heatmaps showing gene expression patterns across all experimental samples of downregulated Naive DEGs (*left*), calculated from Naïve vs 5dpl, 5dpl activated upregulated DEGs (*center*), calculated from 5dpl vs Naïve, and 20dpl upregulated DEGs (*right*), calculated from 20dpl vs 5dpl, selected by top ranking based on adjusted p value. **D)** Dot plots showcasing top marker genes used to annotate both Astro1 subclusters (*top*) and Astro2 subclusters (*bottom*). See Table S1 for a complete list of cluster markers. **E)** Volcano plot showcasing DEGs calculated from a pairwise comparison of all Astro1 subclusters (*purple*) vs Astro2 subclusters (*blue*). Log2FC >1.0, min. pct = 0.2, adjusted p-value <0.05, MAST, Bonferroni correction. **F)** Bar plot showcasing gene set enrichment terms associated with calculated upregulated DEGs between naïve Astro1 astrocytes vs Astro2, organized by −log10(p-value). Bars represent the −log10(p-value) for upregulated DEGs in Astro1 (*blue*) and upregulated DEGs in Astro2 (*orange*). Gene set enrichment was conducted using Metascape. **G)** Feature plot (*left*) and violin plot (*right*) showcasing the expression of Astro1 markers *Slc8a1* and *A2m* (*top*), and Astro2 markers *Slc7a10* and *Gpc5* (*bottom*). Black dot in violin plot represents the median expression. **H)** Scaled heatmaps showing differential gene expression patterns in Astro1 (Astro1, iAstro1a, iAstro1b, DA-Astro, pAstro1) nuclei across all experimental samples. Selected genes were acquired form pairwise DEG testing between 20dpl vs 5dpl. Lipid-related DEGs (*top*) and glutamate signaling-related DEGS (*bottom*). **I)** Heatmap showing the correlation (Pearson) between annotated astrocyte subclusters based on the top 2000 variable genes. The black outline box highlights Astro1 subclusters, while the gray outline box highlights Astro2 subclusters. **J)** Scaled heatmaps showing differential gene expression patterns across grouped nuclei subclusters Astro1 (Astro1), iAstro1 (iAstro1a, iAstro1b, pAstro1), DA-Astro (DA-Astro), Astro2 (Astro2), and iAstro2 (iAstro2a, iAstro2b, iAstro2c, and IFN-R). Selected genes were acquired from gene sets for ECM organization (*top*) and vasculature development (*bottom*). **K)** Venn diagram showcasing the overlap (177 genes) of upregulated DEGs in injury-associated astrocytes calculated from pairwise comparisons of iAstro1 vs Astro1 (516 unique) and iAstro2 vs Astro2 (149 unique). The top 20 upregulated genes unique to iAstro2 (*top, green*) and iAstro1 (*bottom, black*) are shown.

**Figure S5.**
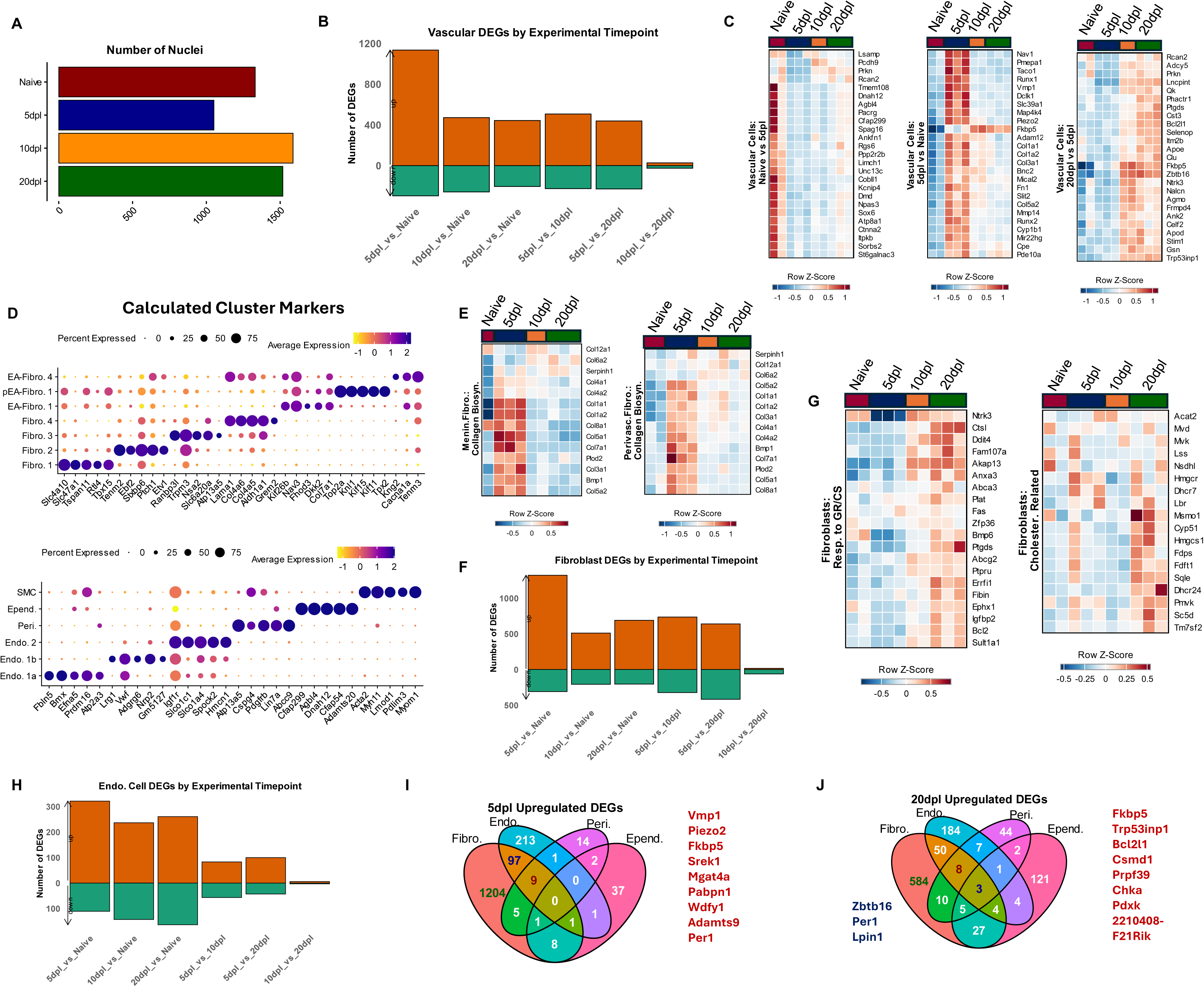
Characterization of vascular and mesenchymal subclusters throughout remyelination. **A)** Bar plot showing the number of vascular and mesenchymal nuclei per timepoint. **B)** Stacked bar plot indicating the number of upregulated (*top, orange*) and downregulated (*bottom, green*) genes calculated in pairwise comparisons in all vascular and mesenchymal nuclei between each experimental timepoint (5dpl, 10dpl, 20dpl, and Naïve). Log2FC >1.0, min. pct = 0.2, adjusted p-value <0.05, MAST, Bonferroni correction. **C)** Scaled heatmaps showing gene expression patterns across all experimental samples of downregulated 5dpl DEGs (*left*), and 5dpl activated upregulated DEGs (*center*), calculated from 5dpl vs Naïve, and 20dpl upregulated DEGs (*right*), calculated from 20dpl vs 5dpl, selected by top ranking based on adjusted p value. **D)** Dot plots showcasing top marker genes used to annotate both fibroblast subclusters (*top*) and endothelial subclusters, pericytes, ependymal cells, and SMCs (*bottom*). See Table S1 for a complete list of cluster markers. **E)** Scaled heatmaps showing differential gene expression patterns of collagen biosynthesis genes in meningeal fibroblasts (*left*) and perivascular fibroblasts (*right*) across all experimental samples. Naïve samples were combined for meningeal fibroblasts as they were lowly detected. **F)** Stacked bar plot indicating the number of upregulated (*top, orange*) and downregulated (*bottom, green*) genes calculated in pairwise comparisons in fibroblast nuclei between each experimental timepoint (5dpl, 10dpl, 20dpl, and Naïve). Log2FC >1.0, min. pct = 0.2, adjusted p-value <0.05, MAST, Bonferroni correction. **G)** Scaled heatmaps showing differential gene expression patterns in all fibroblast nuclei across all experimental samples. Selected genes were acquired from gene sets for response to glucocorticoid and corticosteroid signaling (*left*) and cholesterol related (*right*). **H)** Stacked bar plot indicating the number of upregulated (*top, orange*) and downregulated (*bottom, green*) genes calculated in pairwise comparisons in endothelial cell nuclei between each experimental timepoint (5dpl, 10dpl, 20dpl, and Naïve). Log2FC >1.0, min. pct = 0.2, adjusted p-value <0.05, MAST, Bonferroni correction. **I)** Venn diagram showcasing the overlaps of upregulated 5dpl DEGs in Fibroblasts, Endothelial cells, Pericytes, and Ependymal cells calculated from pairwise comparisons of 5dpl vs Naïve in each broad subpopulation respectively. The upregulated genes shared between fibroblasts, endothelial cells, and pericytes (*red*) are shown. **J)** Venn diagram showcasing the overlaps of upregulated 20dpl DEGs in Fibroblasts, Endothelial cells, Pericytes, and Ependymal cells calculated from pairwise comparisons of 20dpl vs Naïve in each broad subpopulation respectively. The upregulated genes shared between fibroblasts, endothelial cells, and pericytes (*red*) and those shared in all groups (*blue*) are shown.

**Figure S6.**
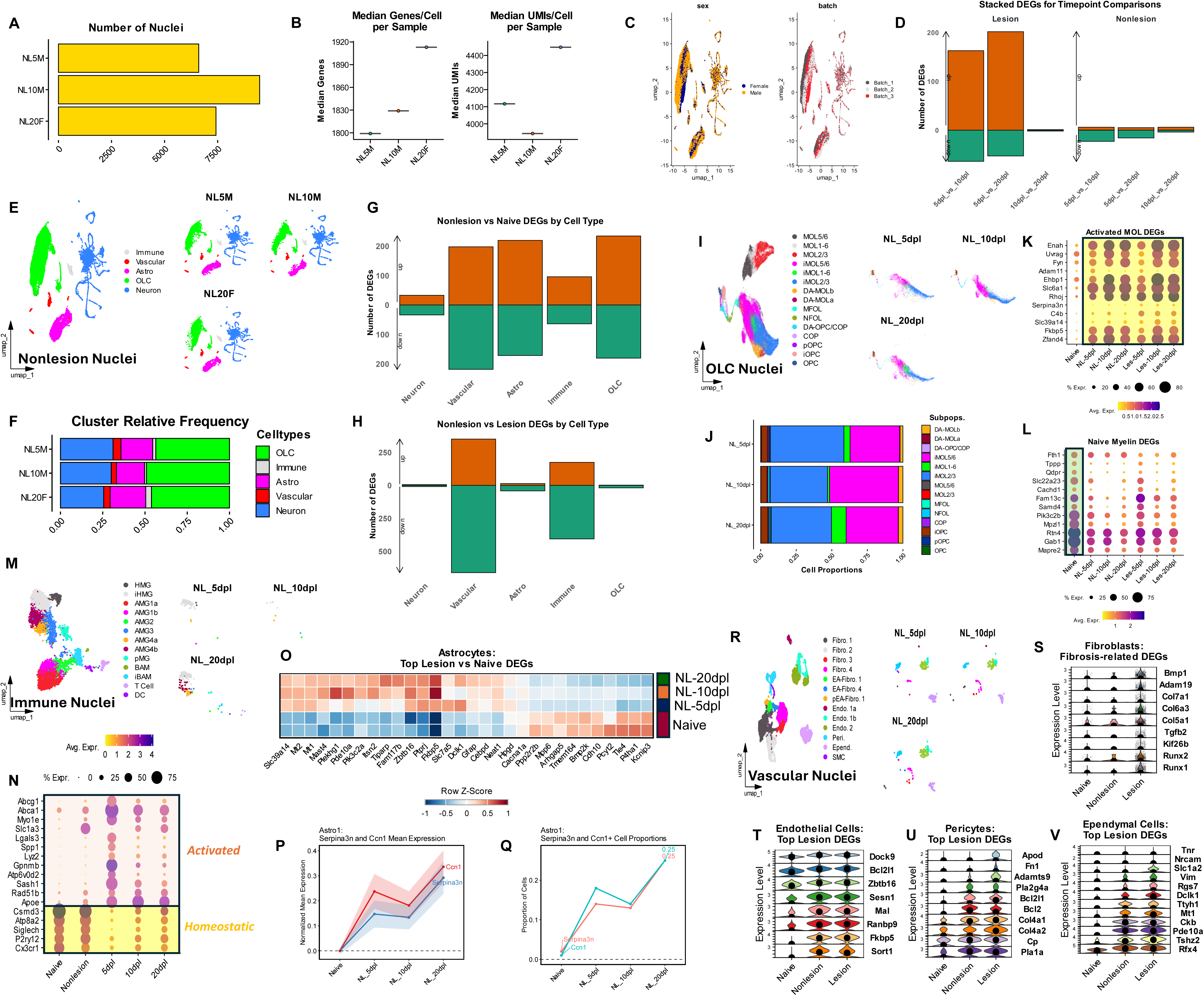
Characterization of adjacent, nonlesioned control tissues throughout remyelination. **A)** Bar plot showing the number of overall nonlesioned sample nuclei per timepoint. **B)** Box and whisker plots showing the median genes/cell per experimental sample (*left*) and median UMIs/cell per experimental sample (*right*) across nonlesioned timepoints. **C)** UMAP projections split (clustered) by sex (*left*) and batch (*right*) indicating together the lack of technical batch effects. **D)** Stacked bar plot indicating the number of upregulated (*top, orange*) and downregulated (*bottom, green*) genes calculated in pairwise comparisons across all nuclei between each experimental timepoint in lesioned samples (*left*) and nonlesioned samples (*right*). Log2FC >1.0, min. pct = 0.2, adjusted p-value <0.05, MAST, Bonferroni correction. **E)** UMAP plot of broadly annotated nonlesioned nuclei (*left*), split by nonlesioned timepoint (*right*). **F)** Bar plot showing the relative frequencies of annotated broad cell subpopulations in each nonlesioned timepoint, highlighting the similarities between them. See Table S2 for the number of nuclei and relative frequencies of all annotated clusters. **G)** Stacked bar plot indicating the number of upregulated (*top, orange*) and downregulated (*bottom, green*) genes calculated in pairwise comparisons of each major cell population between nonlesioned samples and naïve controls. Log2FC >1.0, min. pct = 0.2, adjusted p-value <0.05, MAST, Bonferroni correction. **H)** Stacked bar plot indicating the number of DEGs calculated in pairwise comparisons of each major cell population between nonlesioned and lesioned samples. Brown bar indicates upregulation in nonlesioned samples, and green bar indicates upregulation in lesioned samples. Log2FC >1.0, min. pct = 0.2, adjusted p-value <0.05, MAST, Bonferroni correction. **I)** UMAP plot of annotated OLC nuclei in lesioned, nonlesioned, and naïve samples (*left*), split to highlight the nonlesioned samples by timepoint (*right*). **J)** Bar plot showing the relative frequencies of annotated OLC subpopulations in each nonlesioned timepoint. See Table S2 for the number of nuclei and relative frequencies of all annotated clusters. **K)** Dot plot showcasing top injury-associated/activated genes observed in MOLs, compared between all timepoint conditions in the dataset. **L)** Dot plot showcasing top myelin-related genes observed in naive MOLs, compared between all timepoint conditions in the dataset. **M)** UMAP plot of annotated immune nuclei in lesioned, nonlesioned, and naïve samples (*left*), split to highlight the nonlesioned samples by timepoint (*right*). **N)** Dot plot of myeloid cells showcasing top homeostatic and activated genes used to differentiate activation status in microglial cells across experimental timepoints. **O)** Scaled heatmaps showing gene expression patterns across naïve and nonlesioned samples of astrocyte DEGs calculated in Figure 4, to highlight the activation profiles evident in nonlesioned astrocytes. **P)** Trendlines indicating the normalized mean gene expression of *Serpina3n* (*blue*) and *Ccn1* (*red*) in Astro1 astrocytes across naïve and nonlesioned timepoints, indicating peak levels at 20dpl. **Q)** Trendlines indicating the proportion of *Serpina3n* (*red*) and *Ccn1* (*blue*) positive nuclei in Astro1 astrocytes across naïve and nonlesioned timepoints, indicating peak levels at 20dpl. **R)** UMAP plot of annotated vascular and mesenchymal nuclei in lesioned, nonlesioned, and naïve samples (*left*), split to highlight the nonlesioned samples by timepoint (*right*). **S-V)** Stacked violin plots showing genes upregulated in lesioned samples across naïve, nonlesioned, and lesioned experimental conditions in fibroblasts (**S**), endothelial cells (**T**), pericytes (**U**), and ependymal cells (**V**). Central dot represents median gene expression.

